# Lipid Nanoparticles for the Delivery of CRISPR/Cas9 Machinery to Enable Site-Specific Integration of CFTR and Mutation-Agnostic Disease Rescue

**DOI:** 10.1101/2025.01.22.633938

**Authors:** Ruth A. Foley, Paul G. Ayoub, Vrishti Sinha, Colin Juett, Alicia Sanoyca, Emily C. Duggan, Lindsay E. Lathrop, Priyanka Bhatt, Kevin Coote, Beate Illek, Brigitte N. Gomperts, Donald B. Kohn, Steven J. Jonas

## Abstract

We report the engineering of lipid nanoparticles (LNPs) to transport CRISPR/Cas9 payloads, including double-stranded DNA (dsDNA) donor templates, designed for homology directed repair (HDR)-mediated site-specific insertion of the cystic fibrosis transmembrane conductance regulator (*CFTR*) gene to correct cystic fibrosis (CF) in diseased airway epithelium. We screened various nanoparticle formulations, adjusting ratios of Cas9-encoding mRNA, single guide RNAs (sgRNAs), and dsDNA donor templates to optimize gene editing using human bronchial epithelial cells (16HBE14o-) harboring a CF-causing mutation (G542X). Populations of G542X cells edited *via* LNP delivery of *CFTR* donors achieved 3 – 3.5% gene integration and yielded comparable CFTR protein expression compared to normal 16HBE14o- controls. These edited populations exhibit restoration of CFTR-dependent Cl- current to *ca.* 80% of values measured in normal 16HBE14o- cell monolayers. This LNP platform adds capabilities for transporting large gene editing machinery to airway epithelial cells for genomic integration of entire genes, enabling therapeutic solutions that achieve correction of any CF-causing mutation.

## Introduction

Gene editing approaches, such as clustered regularly interspersed short palindromic repeats (CRISPR)/CRISPR-associated protein 9 (CRISPR/Cas9) technology and its rapidly advancing derivatives, are revolutionizing the treatment landscape for patients with genetic disorders. These tools promise sustained correction of the underlying causes of genetic diseases by manipulating patients’ genomes to knock out pathogenic genes, correct disease-causing mutations, or insert healthy copies of genes (*1–3*). Successful clinical implementation of these genome editing technologies necessitates coordinated intracellular transport of required biomolecular components (*i.e.,* Cas9 nucleases, guide RNAs, and/or homologous donor DNA templates) (*4*).

Lipid nanoparticles (LNPs) have emerged as a promising platform for gene delivery, especially following the widespread success of the Pfizer-BioNTech and Moderna COVID-19 vaccines, which leverage LNPs to deliver mRNA encoding the COVID-19 spike protein (*5, 6*). Prior to the COVID-19 pandemic, LNPs had already demonstrated utility for delivering siRNA (*4, 6, 7*). Since then, the application of new LNP-based delivery solutions for genes and gene-editing components has accelerated, driven by growing interest in alternatives to viral vectors.

One area of particularly rapid development is in gene delivery strategies for targeting therapeutic solutions for cystic fibrosis using LNPs. Cystic fibrosis (CF) is an appealing target for gene therapy, as it stems from mutations in a single gene—the cystic fibrosis transmembrane conductance regulator (*CFTR*) gene—that result in impaired ion transport across epithelia (*8–10*). In the airway epithelium, dysfunctional ion transport results in formation of dehydrated, sticky mucus and impaired mucociliary clearance (*10–12*). Patients suffer from chronic infections that lead to progressive pulmonary scarring and eventual respiratory failure. While many patients can be managed successfully with modulator therapies that augment the function of the CFTR protein, approximately 10% of patients have mutations that are not responsive to these interventions (*13*). These mutations are a particular health disparity concern, as they are overrepresented in non-white patient populations (*13*). For patients with untreatable mutations, gene therapies promise definitive treatment.

Durable gene editing of airway basal stem cells (ABSCs) to express functional CFTR would result in long-lived alleviation of respiratory symptoms for patients harboring any of the *ca.* 2000 CF-causing mutations (*8–10*). Recently, LNPs have been demonstrated to achieve robust gene editing in CF models using a CRISPR/Cas9 approach that leveraged single-stranded oligonucleotide (ssODN) donor templates (*14*). These short (*ca.* 200 nucleotides) homology directed repair (HDR) templates achieved robust correction of multiple CF-causing mutations in both primary human bronchial epithelial (HBE) cell models and in mice (*14*). Further, editing approaches that do not rely on the formation of double-stranded breaks (DSBs), such as base editors, have recently shown promise for correcting specific individual CF-associated mutations (*15*).

Several CRISPR-based strategies (*e.g.,* Cas9-mediated insertion *via* HDR or homology-independent targeted integration (HITI); CRISPR-associated transposases or CASTs; CRISPR-associated recombinases/integrases) rely on large DNA donor templates for gene editing (*16, 17*). These systems offer a few distinct advantages over mutation-specific correction using smaller ssODN donor templates or *via* base and prime editing approaches. First, insertion of long sequences offers a mutation-agnostic platform that does not require mutation-specific customization, reducing cost and enhancing scalability of the therapy. Second, insertion of entire corrected gene offers the flexibility to utilize codon-optimized and/or gain-of-function variants of the donor cassette, increasing protein expression and/or function and potentially overcoming limitations to delivery and editing efficiencies (*18–22*). However, the packaging capacity of synthetic nanocarriers like LNPs for these large cargoes is still poorly understood, and LNPs have not yet been applied effectively to deliver linear double stranded DNA (ldsDNA).

Here, we report the design and validation of a LNP platform that simultaneously packages mRNA transcripts encoding Cas9, single guide RNAs (sgRNAs), and large ldsDNA donor template cargoes (Fig. 1). These reagents have been developed by our group to enable site-specific integration of a codon-optimized donor template encoding the 5kb *CFTR* coding sequence with 50 bp homology arms within the 5’ untranslated region (UTR) of the endogenous *CFTR* gene with no detectable off-target activity (*23*). This site-specific insertion strategy leaves intact the endogenous *CFTR* promoter and introns, allowing native intracellular machinery to regulate transcription of the integrated sequence. Our work serves as proof-of-concept that LNPs can successfully deliver large (>5kb) ldsDNA transcripts and offer an exciting delivery modality for gene editing reagents that have not yet been applied in conjunction with this nanocarrier platform.

**Figure 1.**
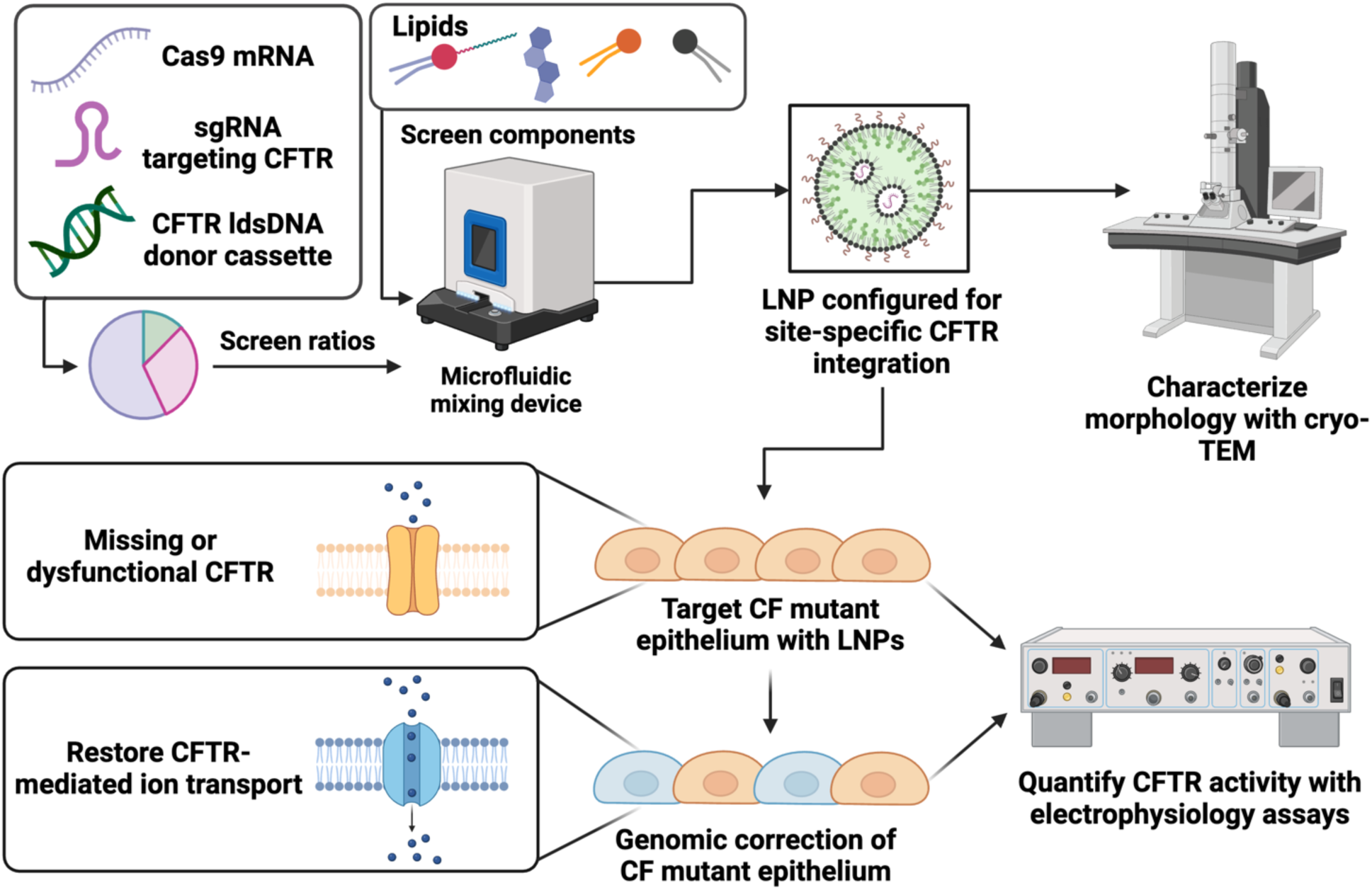
Schematic overview of study design and workflow. Lipid nanoparticles (LNPs) were first optimized by screening individual lipid components for efficient delivery to human bronchial epithelial cells then further configured to deliver Cas9 mRNA, single guide RNA (sgRNA) targeting the cystic fibrosis transmembrane conductance regulator (*CFTR*) gene, and linear double-stranded DNA (ldsDNA) donor cassettes encoding *CFTR.* The encapsulated ratios of gene-editing cargoes were systematically screened to maximize site-specific genomic integration of the ldsDNA donor cassette. Particles were characterized using cryogenic transmission electron microscopy (cryo-TEM) and applied to CF mutant epithelial cells to achieve *CFTR* gene correction. CFTR-dependent ion transport of edited cell populations was quantified using transepithelial current clamp (TECC) assays.

## Results

### Selection of lipid nanoparticle constituents

Traditional LNP formulations comprise an ionizable lipid, a helper phospholipid, a sterol, and a lipid conjugated to poly(ethylene glycol) (PEG). The exact identities and molar ratios of each of these constituents must be optimized according to application, since LNP self-assembly depends on the nature of the encapsulated cargo, and the biology of LNP uptake and endosomal trafficking varies according to cell type (*24, 25*). With these considerations in mind, we aimed to establish a LNP formulation optimized for nucleic acid delivery to a disease-relevant bronchial epithelial cell line (16HBE14o-). Initial formulations were composed of 50 mol% ionizable lipid (IL-A), 38.5 mol% sterol (sterol-A), 10 mol% 1,2-dioleoyl-sn-glycero-3-phosphoethanolamine (DOPE) and 1.5 mol% lipid-PEG (LP-A). Different lipid components were systematically substituted into this base formula to screen for their ability to package and deliver reporter mRNA cargoes encoding green fluorescent protein (GFP) to 16HBE14o- cells.

We first tested commercially available ionizable lipids containing degradable ester groups including lipids IL-B and IL-C against IL-A as candidate LNP components. Each of these ionizable lipids were compared for their ability to form small, uniform LNPs with high (>80%) encapsulation efficiencies as measured by the RiboGreen assay (Thermo Fisher) and their delivery capabilities. All ionizable lipids produced LNPs with similar sizes and encapsulation efficiencies (Fig. 2). LNPs prepared using IL-A achieved GFP expression *ca.* 7-8% higher in targeted 16HBE14o- cells 24 h after LNP treatment as measured by flow cytometry than LNPs made with IL-B (*p* = 0.0003) or IL-C (*p* = 0.0007) (Fig. 3E). All LNPs preserved 85 – 95% viability across the three ionizable lipids (n.s.) (Fig. 3F). Based on these findings, we selected IL-A as the optimal ionizable lipid candidate.

**Figure 2.**
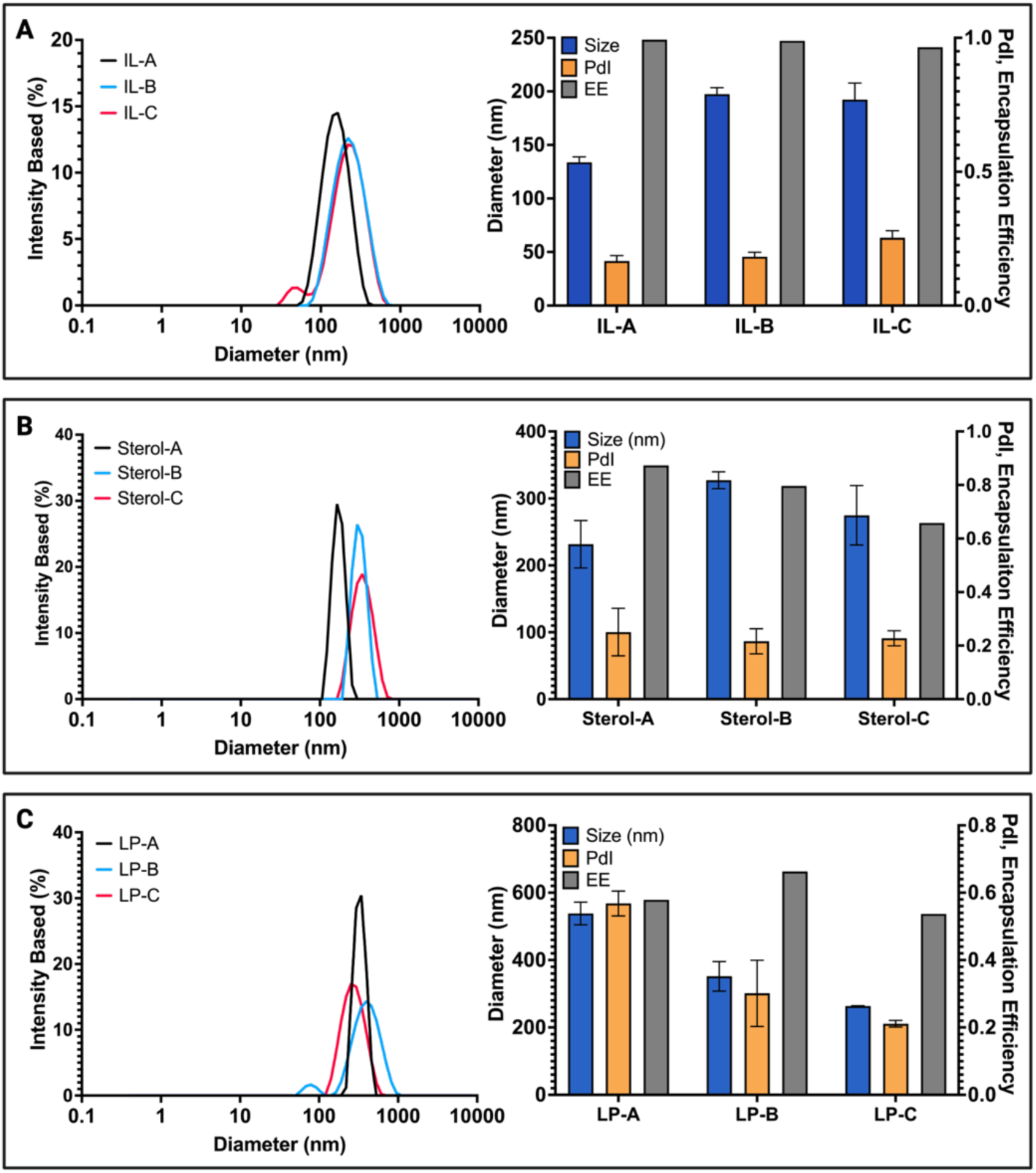
Characterization of candidate lipid nanoparticles (LNPs). (**A**) Dynamic light scattering (DLS) and encapsulation data depicting the size distributions (left) and diameter, polydispersity (PdI), and encapsulation efficiencies (right) of LNPs packaged with enhanced GFP mRNA and containing either lipids IL-A, IL-B, or IL-C. (**B**) DLS and encapsulation data depicting the size distributions (left) and diameter, PdI, and encapsulation efficiencies (right) of LNPs packaged with enhanced GFP mRNA and containing either sterol-A, sterol-B, or sterol-C. (**C**) DLS and encapsulation data depicting the size distributions (left) and diameter, PdI and encapsulation efficiencies (right) of LNPs packaged with GFP mRNA and containing either lipid-pegs LP-A, LP-B, or LP-C.

**Figure 3.**
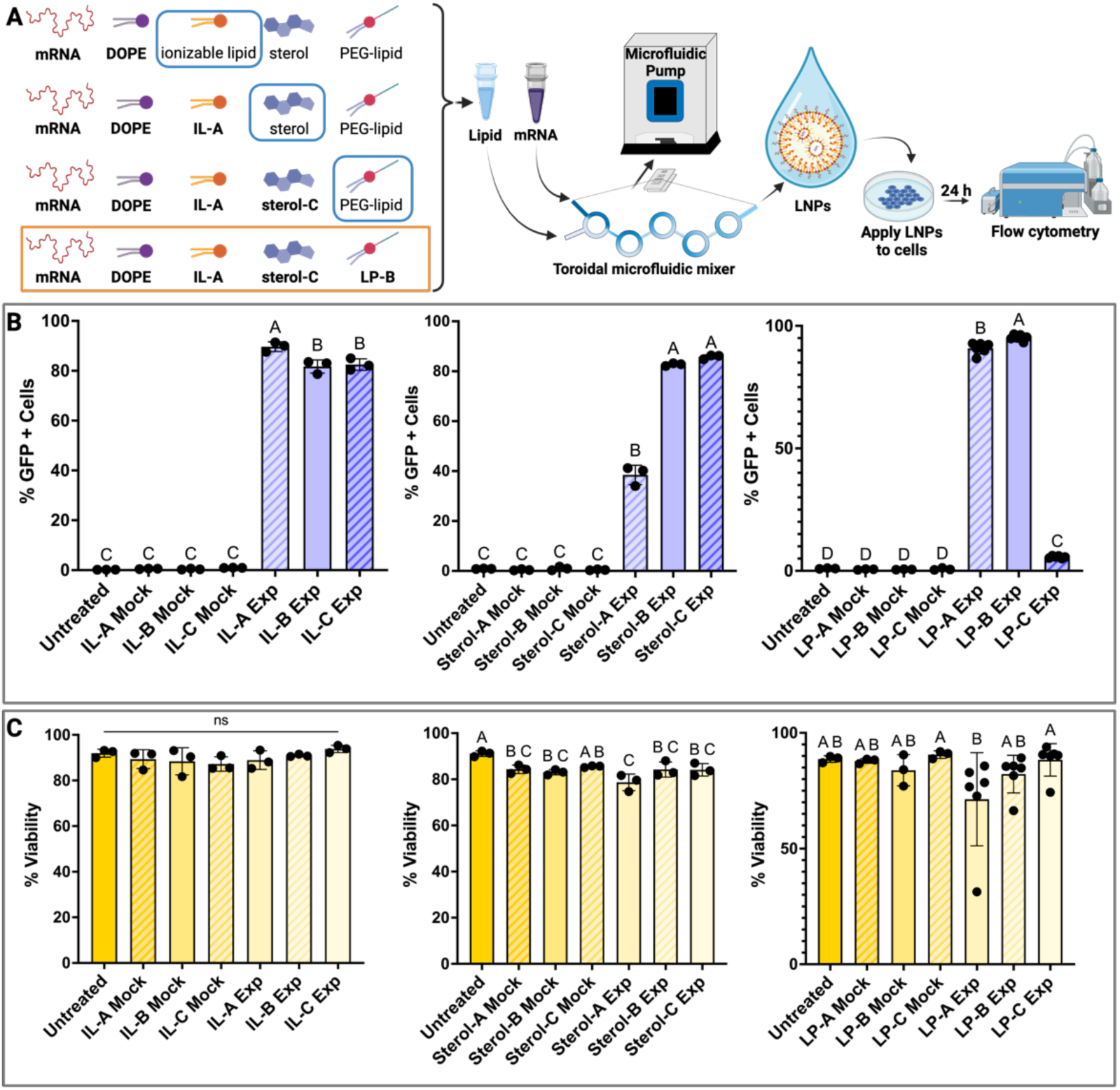
Evaluation of candidate lipid nanoparticle (LNP) formulations for their transfection capabilities. (**A**) Schematic depicting stepwise selection of lipid nanoparticle (LNP) constituents for optimized delivery of mRNA to a human bronchial epithelial cell line (16HBE14o-) and workflow for synthesis of LNPs using a bifurcating toroidal microfluidic mixer and screening for LNP-mediated delivery of reporter mRNA constructs encoding green fluorescent protein (GFP) to 16HBE14o- cells. Optimized formulation includes 50 mol% IL-A, 38.5 mol% sterol-C, 10 mol% 1,2-dioleoyl-sn-glycero-3- phosphoethanolamine (DOPE), and 1.5 mol%, LP-B. (**B**) Flow cytometry data depicting percentage GFP-expressing 16HBE14o- cells 24 h after treatment with LNPs containing different (left) ionizable lipids, (center) sterols, and (right) PEG-lipids. (**C**) Cell viability indicated by 7-AAD staining and measured by flow cytometry in 16HBE14o- cells 24 h after treatment with LNPs containing different (left) ionizable lipids, (center) sterols, and (right) PEG-lipids. One-way analyses of variance (ANOVA) or Kruskal-Wallis tests of variance were performed, with the threshold of statistical significance set at *P <* 0.05. Letters A-D represent a compact letter display of statistics wherein differences between groups labeled with the same letter are not statistically significant.

We further investigated whether delivery of mRNA-based cargoes could be enhanced by substituting sterol-A with either sterol-B or sterol-C. Nanoparticles containing sterol-A tended toward higher encapsulation efficiencies and smaller sizes (Fig. 2B). We observed sterol-C LNPs to achieve *ca.* 5-10% higher GFP expression than particles containing sterol-B (n.s.) and 30 – 40% higher than particles that incorporated sterol-A (*p* < 0.0001, Fig. 3E).

Finally, the impact of the structure and length of the lipid-PEG component was probed to determine its effect on LNP size, encapsulation efficiency, and transfection performance. We directly compared encapsulation and delivery of GFP-encoding mRNA constructs in LNPs containing either PL-A, PL-B, or PL-C. Regardless of PEG-lipid, all LNP formulations tested exhibited similar sizes and encapsulation efficiencies (Fig. 2C). LNPs containing LP-B achieved 4-5% greater GFP expression compared to LNPs containing LP-A (*p* < 0.0001) and *ca.* 90% higher expression compared to LP-C LNPs (*p* < 0.0001) (Fig. 3E). While LP-A LNPs appeared to preserve cell viability better than LNPs made using LP-C (*ca.* 16%, *p* = 0.0364), there was no significant difference in viability between LNPs containing LP-A and LP-B (Fig. 3F).

LNPs comprised entirely of optimized lipid building blocks achieve transfection efficiencies of 95% (Fig. 3E) and preserved >80% cell viability (Fig. 3F). Specifically, this formula includes 50 mol% IL-A, 38.5 mol% sterol-C, 10 mol% DOPE, and 1.5 mol% LP-B.

### Screening ratios of gene-editing cargoes

While the optimized LNP formulation successfully demonstrated delivery of small model GFP transcripts efficiently to 16HBE14o- cells, LNP-mediated gene editing relies on the delivery of multiple cargoes at once. Specifically, our HDR-mediated site-specific insertion approach requires the intracellular delivery of a mRNA transcript encoding a Cas9 endonuclease, a sgRNA designed to target a specific genomic locus, and a donor DNA construct that serves as a template for the sequence to be inserted. We designed a set of experiments to screen the impact of relative ratios of these components on the gene-editing capabilities of our LNPs. For these studies, we generated a 16HBE14o- model reporter line stably expressing blue fluorescent protein (BFP) according to a protocol from Corn and colleagues (Fig. 4A) (*26*). LNPs were loaded with Cas9 mRNA, a sgRNA targeting the BFP locus, and a single-stranded oligonucleotide (ssODN) donor designed to introduce a three-nucleotide mutation (targeting H66) to convert BFP to GFP *via* HDR. A loss of cell fluorescence indicates non-homologous end joining (NHEJ). This reporter system offers a convenient tool for rapidly assessing overall editing efficiencies and HDR:NHEJ ratios *via* flow cytometry. We systematically screened ratios of the mRNA, sgRNA, and ssODN cargoes to determine the conditions that maximize HDR-mediated editing. Initially, the ssODN:mRNA ratio was fixed at 3:1 w/w and the sgRNA:mRNA ratio was systematically titrated from 0.8:1 w/w to 2:1 w/w. For all ratios, LNPs induced 10 – 20% HDR and 25 – 40% overall editing, with no significant differences reproducibly observed between any sgRNA:mRNA ratios (Fig. 4B). Cell viability for all conditions was maintained at *ca.* 80 – 90% (Supplementary Materials Fig. S3). Biological replicates of this experiment show 4-6% greater incidence of HDR for LNPs loaded at a sgRNA:mRNA ratio of 1.2:1 w/w as compared to 0.8:1 w/w (*p* = 0.0002) and 2:1 w/w (*p* = 0.0101), with no significant difference compared to 1.6:1 w/w (Supplementary Materials Fig. S3). We therefore selected a sgRNA:mRNA ratio of 1.2:1 w/w for use in subsequent experiments.

**Figure 4.**
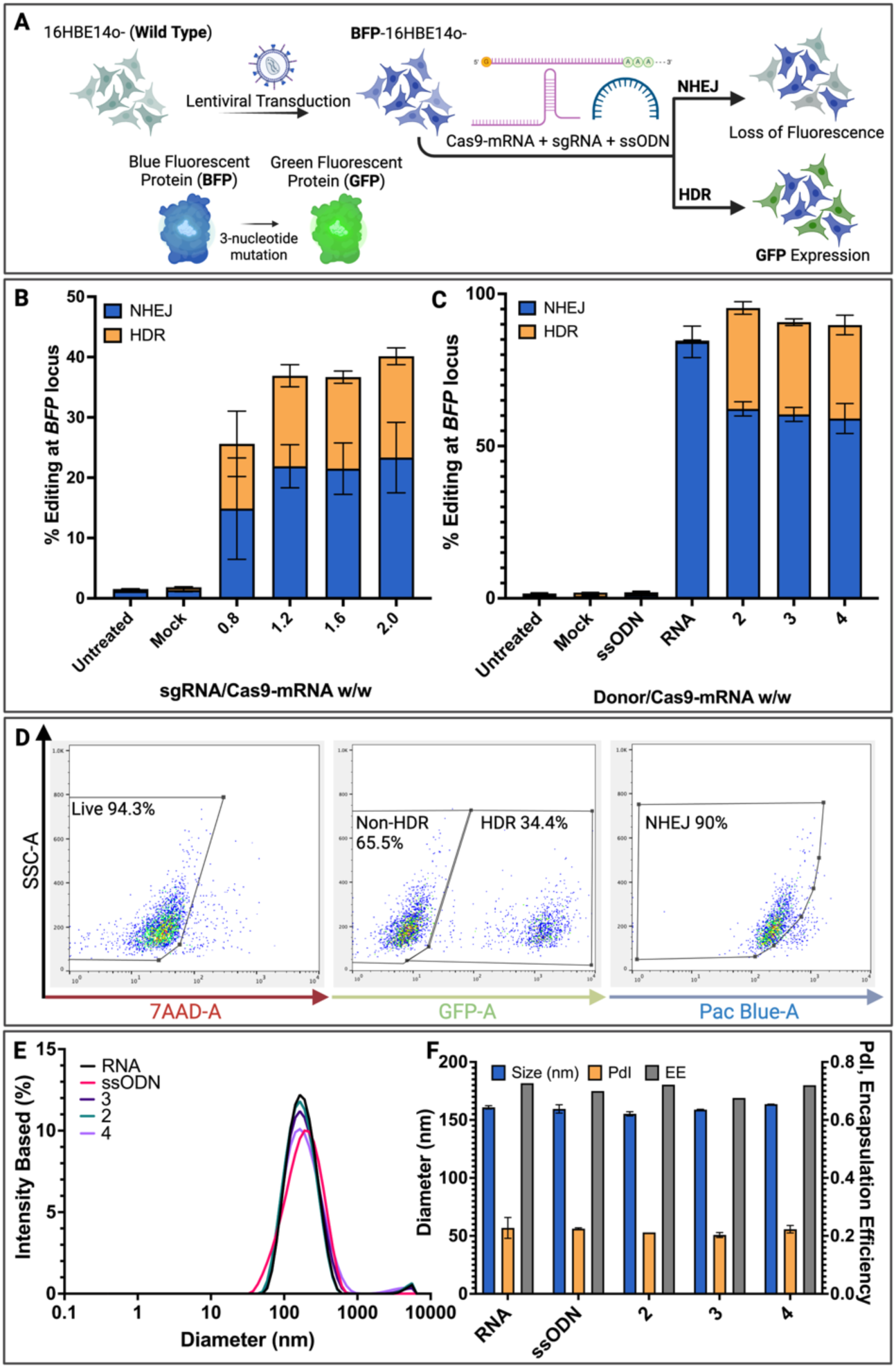
Optimization of a lipid nanoparticle (LNP) formulation for gene editing. **(A)** Schematic illustrating the blue fluorescent protein (BFP)-expressing 16HBE14o- reporter line used as a tool to assess capabilities of LNPs to deliver gene editing cargos including mRNA constructs encoding Cas9, single guide RNA (sgRNA), and a single-stranded oligonucleotide (ssODN) donor. Loss of blue fluorescence indicates non-homologous end-joining (NHEJ), and expression of green fluorescent protein (GFP) indicates homology-directed repair (HDR)-mediated editing. (**B**) Flow cytometry data depicting NHEJ and HDR efficiencies in BFP-expressing 16HBE14o- cells 5 days after treatment with LNPs loaded with mRNA transcripts encoding Cas9, sgRNA, and an ssODN targeting the BFP gene. The ratio of ssODN to mRNA was held constant at 3:1 w/w, with the ratio of sgRNA to mRNA titrated from 0.8:1 w/w to 2:1 w/w. (**C**) Flow cytometry data depicting NHEJ and HDR efficiencies in BFP-expressing 16HBE14o- cells 5 days after treatment with LNPs loaded with mRNA transcripts encoding Cas9, sgRNA, and ssODNs targeting the BFP gene. The ratio of sgRNA to mRNA was held constant at 1.2:1 w/w, with the ratio of ssODN to mRNA titrated from 2:1 w/w to 4:1 w/w. (**D**) Representative flow cytometry plots depicting gating strategies for viability staining (left, 7AAD), HDR (middle, GFP), and NHEJ (right, Pac-Blue). (**E**) Dynamic light scattering (DLS) data depicting the size distributions of LNPs loaded with various ssODN:mRNA w/w ratios. (**F**) DLS data depicting the Z-average diameter and encapsulation efficiencies as measured *via* the RiboGreen assay (Thermo Fisher) of LNPs loaded with various ssODN:mRNA w/w ratios.

Next, the ssODN:mRNA ratio was similarly titrated from 2 to 4 w/w while maintaining the sgRNA:mRNA ratio constant at 1.2:1 w/w. For all ratios, LNPs induced 30 – 35% HDR and 90 – 95% overall editing, with no significant differences reproducibly observed between any ssODN:mRNA ratios (Fig. 4C). Viability for all conditions was maintained at *ca.* 80 – 95% (Supplementary Materials Fig. S3). Biological replicates of this experiment show 3% greater incidence of HDR for LNPs loaded at a ssODN:mRNA ratio of 3:1 w/w as compared to 2:1 w/w (*p* = 0.0031), with no significant difference compared to 4:1 w/w (Supplementary Materials Fig. S3). Based on these findings a ssODN:mRNA ratio of 3:1 w/w was selected for use in subsequent experiments.

### Optimizing LNP formulation to accommodate ldsDNA donor cassettes

Further experiments were performed to determine whether the LNPs tailored for gene editing of *BFP* in 16HBE14o- could be applied toward delivering ldsDNA donor templates encoding for whole genes. As a reporter for successful site-specific gene insertion, a *mCitrine* ldsDNA construct was designed for integration at the 5’UTR of the endogenous *CFTR* gene (*23*). Differences in physicochemical properties of ldsDNA compared to ssODNs were hypothesized to require re-optimization of LNP constituent ratios. Integration efficiencies in 16HBE14o- cells were compared between LNP formulations packaging the Cas9-mRNA, sgRNA, and ldsDNA donor cargoes separately or together within one LNP formulation. Initially, LNPs were prepared at a nitrogen/phosphate (N/P) ratio of 7. Figure 5A compares particle mixtures separating the ldsDNA into a distinct LNP (sepLNPs) and formulations that combine all three gene-editing components into a single LNP (togLNPs). Cargo encapsulation efficiencies were comparable (*ca.* 80%) between togLNPs and sepLNPs containing RNA only (RsepLNPs), and lower (62.6%) for sepLNPs containing DNA only (DsepLNPs) (Fig. 5B). Both togLNPs and RsepLNPs exhibited small (*ca.* 150 nm) diameters and low (*ca.* 0.2) polydispersity indices (PdIs), whereas DsepLNPs were slightly larger (*ca.* 200 nm) with a PDI of 0.34 (Fig. 5B,C). To enhance HDR-mediated integration by downregulating the competing repair processes of non-homologous end joining (NHEJ) and microhomology-mediated end joining (MMEJ), LNPs were co-administered with DNA-dependent protein kinase inhibitor AZD7648 and polymerase-*θ* inhibitor ART558 (*27–29*). Under these conditions, togLNPs achieved integration efficiencies of *ca.* 14%, compared to *ca.* 10% achieved by sepLNPs (*p* = 0.0082) (Fig. 5D,E). When administered without drug, togLNPs achieved 36% greater integration efficiencies than sepLNPs (*p* = 0.0452) (Fig. 5E).

**Figure 5.**
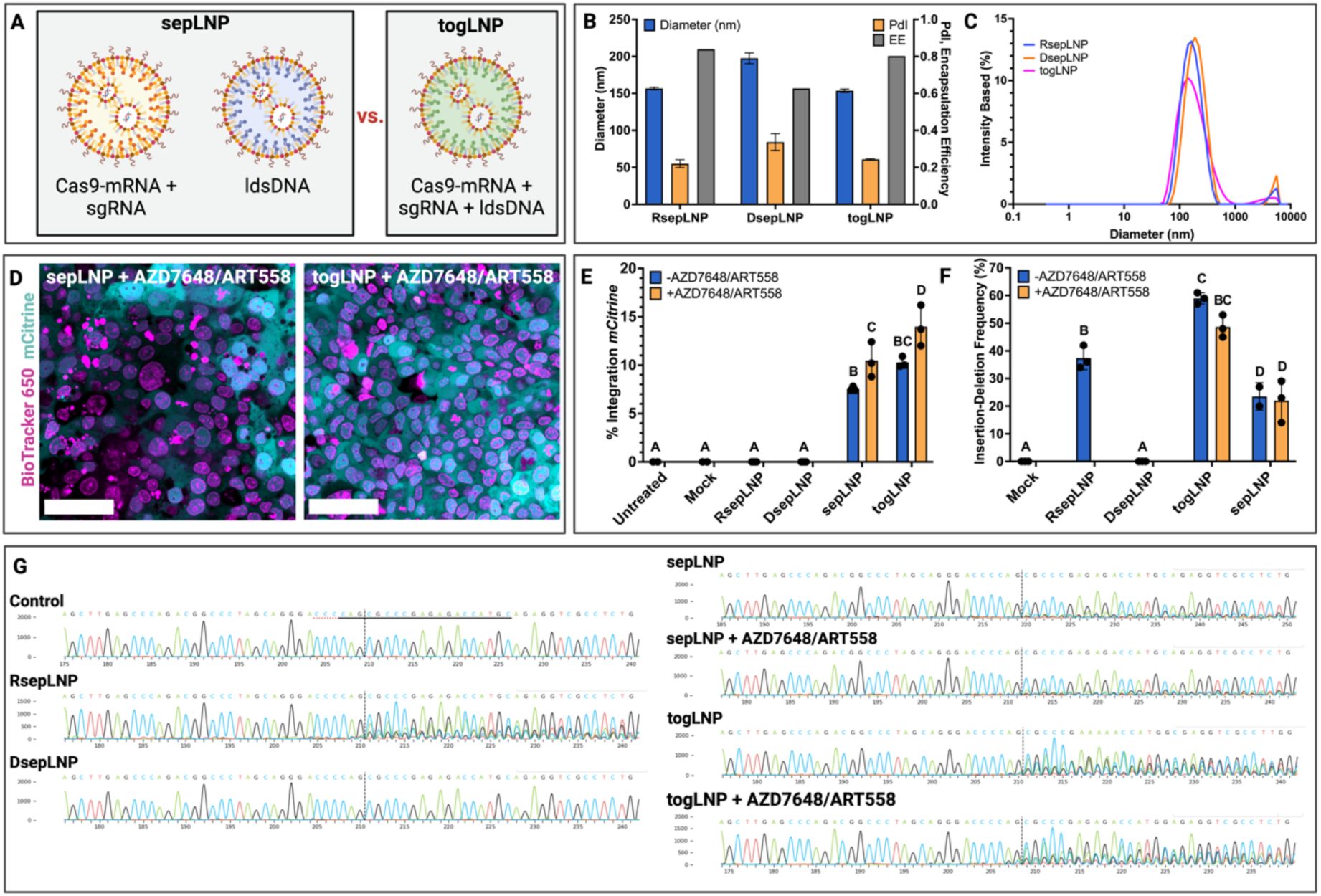
Evaluation of optimized lipid nanoparticles (LNPs) for linear double-stranded DNA (ldsDNA). (**A**) Schematic comparing lipid nanoparticles (LNPs) packaged with mRNA encoding Cas9 (Cas9-mRNA) together with a single-guide RNA (sgRNA) and/or a ldsDNA donor cassette. Particle mixtures separating the ldsDNA into distinct LNPs are referred to as sepLNPs; formulations combining all three gene-editing cargoes into one formulation are referred to as togLNPs. (**B**) Dynamic light scattering (DLS) data depicting the Z-average diameter of RNA-only sepLNPs (RsepLNPs), DNA-only sepLNPs (DsepLNPs), and togLNPs, and encapsulation efficiencies measured *via* RiboGreen assay (ThermoFisher). (**C**) DLS data depicting intensity-based size distributions of RsepLNPs, DsepLNPs, and togLNPs. (**D**) Confocal microscopy images depicting 16HBE14o- cells 5 days after treatment with sepLNPs or togLNPs. Cell nuclei were counterstained with BioTracker 650 (Millipore Sigma). Scale bars = 50 µm. (**E**) Droplet digital PCR (ddPCR) analysis reflecting site-specific integration of an mCitrine reporter cassette within the 5’ untranslated region (UTR) of the endogenous cystic fibrosis transmembrane conductance regulator (*CFTR*) gene in 16HBE14o- cells 48 h after treatment with sepLNPs or togLNPs. (**F**) Inference of CRISPR Edits (ICE) analysis data and (**G**) nucleotide frequency traces depicting rate of insertion-deletions detected within the 5’ UTR of endogenous *CFTR* in 16HBE14o- cells 48 h after treatment with sepLNPs or togLNPs. Cells were incubated overnight with LNPs with or without drugs AZD7648 and ART558. One-way analyses of variance (ANOVA) were performed, with the threshold of statistical significance set at *p <* 0.05.

Integration efficiencies of *mCitrine* were comparable to our results delivering these reagents as Cas9 ribonucleoprotein complexes (RNPs) *via* electroporation (*23*). Furthermore, insertion-deletion (indel) frequencies ranged from 20-25% in samples treated with sepLNPs to >60% in samples treated with togLNPs, indicating overall delivery efficiencies of gene editing cargoes far surpassed integration efficiencies (Fig. 5F,G). These findings prompted additional studies replacing the *mCitrine* reporter construct with a codon-optimized *CFTR-* encoding ldsDNA construct, which we showed in parallel to achieve up to *ca.* 55% restoration of wild-type *CFTR-* dependent ion current at low (1-2%) levels of integration in 16HBEgeG542X cells harboring the G542X premature stop codon mutation (*23*). This mutation was selected as a model for our studies because it is responsible for causing severe CF in roughly 4.5% of patients who are not treatable with traditional CFTR modulator medications (*30*). The *CFTR* construct is nearly five times the size of the *mCitrine* reporter, which required additional optimization of the LNP formulation. The ldsDNA:mRNA ratio was reevaluated, and 2:1, 3:1, and 4:1 w/w were screened and compared (Fig. 6A). A ldsDNA:mRNA ratio of 3:1 was found to achieve greater (up to 3.5%) integration compared to ratios of 2:1 or 4:1, which produced integration efficiencies of <2% (Fig. 6B). No significant differences in cell survival were observed across experimental conditions, with all conditions maintaining 70 – 90% survival relative to the untreated controls (Fig. 6E). A 3:1 w/w ratio of ldsDNA:mRNA was chosen for further experiments. Similarly, editing capabilities were compared between LNPs exhibiting N/P ratios of 5, 7, and 9 (Fig. 6A). Packaging and delivery efficiencies of LNPs are partially dependent on N/P charge ratios that influence the strength of electrostatic interactions between ionizable lipid amines and negatively charged nucleic acid phosphates. When administered with drugs AZD7648 and ART558, LNPs exhibiting a N/P of 9 achieved higher integration efficiencies compared to N/Ps of 5 or 7 with comparable cell survival across all conditions (Fig. 6C,F). To ensure the optimal N/P ratio was captured within the ranges tested, LNPs with N/Ps of 11 and 13 were also evaluated. Integration efficiencies were comparable (*ca.* 1 – 1.5%) between N/Ps of 9, 11, and 13, with particles exhibiting a N/P of 11 achieving the highest integration efficiencies (Fig. 6D). Further, cell survival in samples treated with N/P 9 formulations was lower than samples treated with N/P 11, and significantly lower than samples treated with N/P 13 formulations (*p =* 0.0019, Fig. 6G). For subsequent LNP dosage optimization and functional assays, a N/P of 11 was selected.

**Figure 6.**
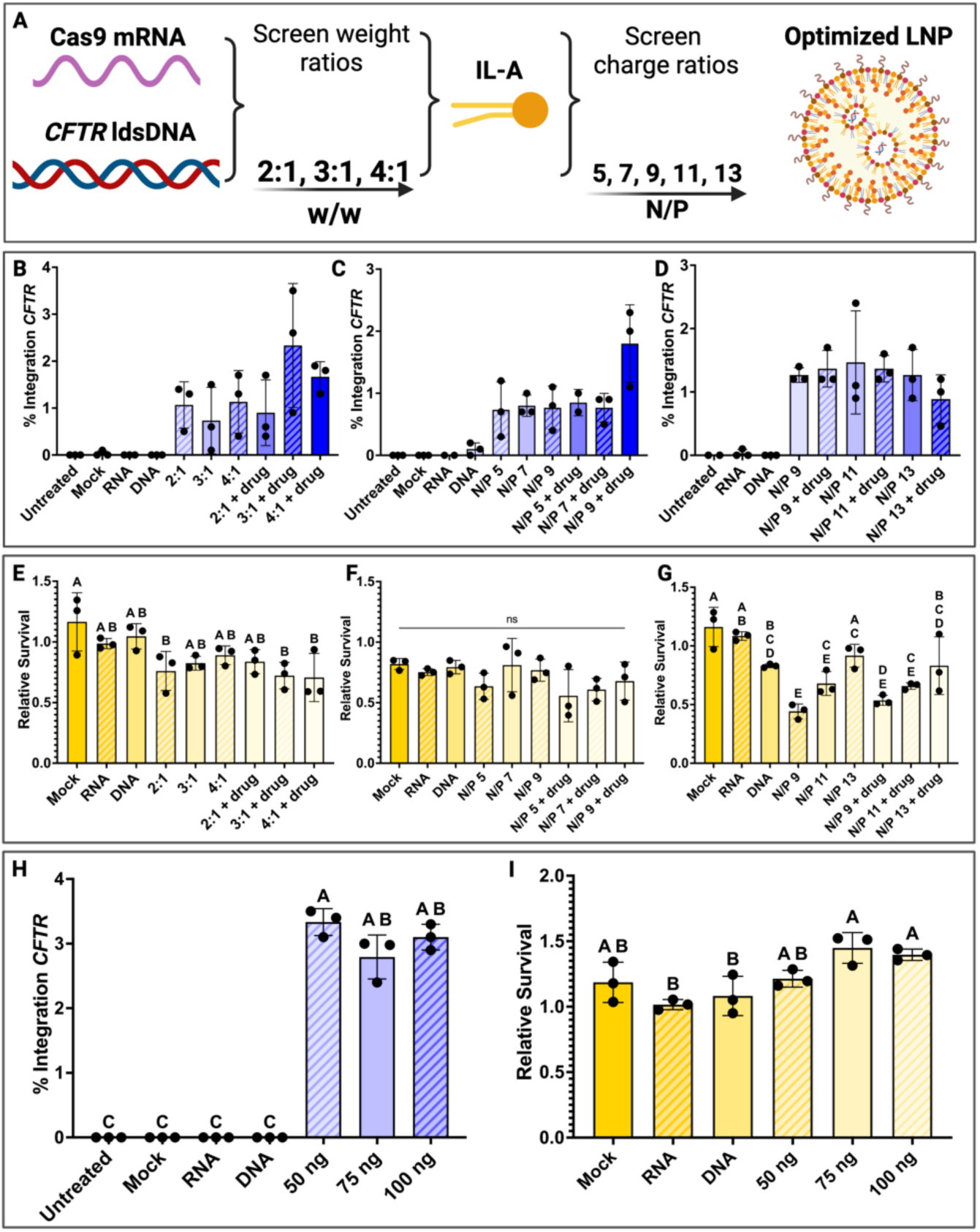
Optimization of lipid nanoparticle (LNP) composition for delivery of functional CFTR-encoding constructs. (**A**) Schematic overview of LNP optimization approach for maximal integration of donor CFTR-encoding linear double-stranded DNA (ldsDNA) constructs. (**B**-**D**) Droplet digital PCR (ddPCR) data reporting integration efficiencies of CFTR ldsDNA constructs co-encapsulated into LNPs with mRNA encoding Cas9 and single guide RNA (sgRNA) targeting *CFTR* at (left) ldsDNA:mRNA w/w ratios of 2:1, 3:1, and 4:1 and (center, right) nitrogen:phosphate (N/P) ratios of 5, 7, 9, 11, or 13. (**E-G**) Metabolic tetrazolium salt (MTS) assay data reporting cell survival relative to untreated cells 24 h after treatment with LNPs loaded with CFTR ldsDNA constructs, mRNA encoding Cas9, and sgRNA targeting *CFTR* at (left) ldsDNA:mRNA w/w ratios of 2:1, 3:1, and 4:1 and (center, right) nitrogen:phosphate (N/P) ratios of 5, 7, 9, 11, or 13. (**H**) Integration efficiencies of CFTR ldsDNA constructs co-encapsulated into LNPs with mRNA encoding Cas9 and sgRNA targeting CFTR at a ldsDNA:mRNA w/w of 3:1 and N/P of 11 at a total dose of 50, 75, or 100 ng mRNA per 20,000 cells. (**I**) MTS assay data reporting cell survival relative to untreated cells 24 h after treatment with LNPs loaded with CFTR ldsDNA constructs, mRNA encoding Cas9, and sgRNA targeting *CFTR* dosed at 50, 75, or 100 ng mRNA per 20,000 cells. Cells were treated with or without drugs AZD7648 and ART558. One-way analyses of variance (ANOVA) were performed, with the threshold of statistical significance set at *p <* 0.05. Letters A-E represent a compact letter display of statistics wherein differences between groups labeled with the same letter are not statistically significant.

The dose of optimized LNPs was titrated from 50 ng mRNA to 100 ng mRNA. All conditions achieved *ca.* 3% integration in the presence of AZD7648 and ART558, with no significant differences between doses (Fig. 6H). The 50 ng condition achieved the highest integration efficiencies with a mean of 3.3% and a maximum of 3.5%, and no significant differences in relative cell survival (Fig. 6H,I).

### LNP-mediated site-specific integration restores CFTR function in G542X-mutant human bronchial epithelial cells

To test whether AZD7648 and ART558 are necessary to achieve comparable integration efficiencies of CFTR ldsDNA, G542X cells were treated with LNPs both with and without these drugs. Neither editing efficiencies nor viabilities were significantly different between samples treated with AZD7648/ART558 (mean integration efficiency = 3.44%) and samples not treated with AZD7648/ART558 (mean integration efficiency = 2.85%, Fig. 7B,C). To evaluate whether editing efficiencies of <4% could confer sufficient *CFTR* expression and function, cells were expanded to conduct Western blot, Ussing chamber, and transepithelial current clamp (TECC) assays (Fig. 7A). Western blot analysis revealed 83% - 123% protein expression in bulk-edited G542X cells (henceforth referred to as G542X-CFTR) compared to normal 16HBE14o- controls (Fig. 7D,E), suggesting that <4% allelic correction can achieve therapeutically relevant CFTR expression levels. Functional assays were performed in our lab using a 2-channel Ussing chamber setup and independently verified by the electrophysiology group at the Cystic Fibrosis Foundation Therapeutics Laboratory (CFFT) using a 24-channel TECC system. Monolayers of G542x-CFTR cells exhibited CFTR chloride currents that were restored to *ca.* 80% - 160% of currents measured in normal 16HBE14o- cell monolayers (Fig. 7F, Supplementary Materials Fig. S6, *p =* 0.0045). Transepithelial electrical resistance (TEER) measurements were >800 Ωxcm^2^ for all samples assayed using the TECC system and consistent across replicates (Supplementary Materials Fig. S7). To our knowledge, this is the first demonstration of ldsDNA cassettes of this size being packaged and delivered intracellularly to achieve phenotypic rescue of a dysfunctional gene *via* a LNP platform.

**Figure 7.**
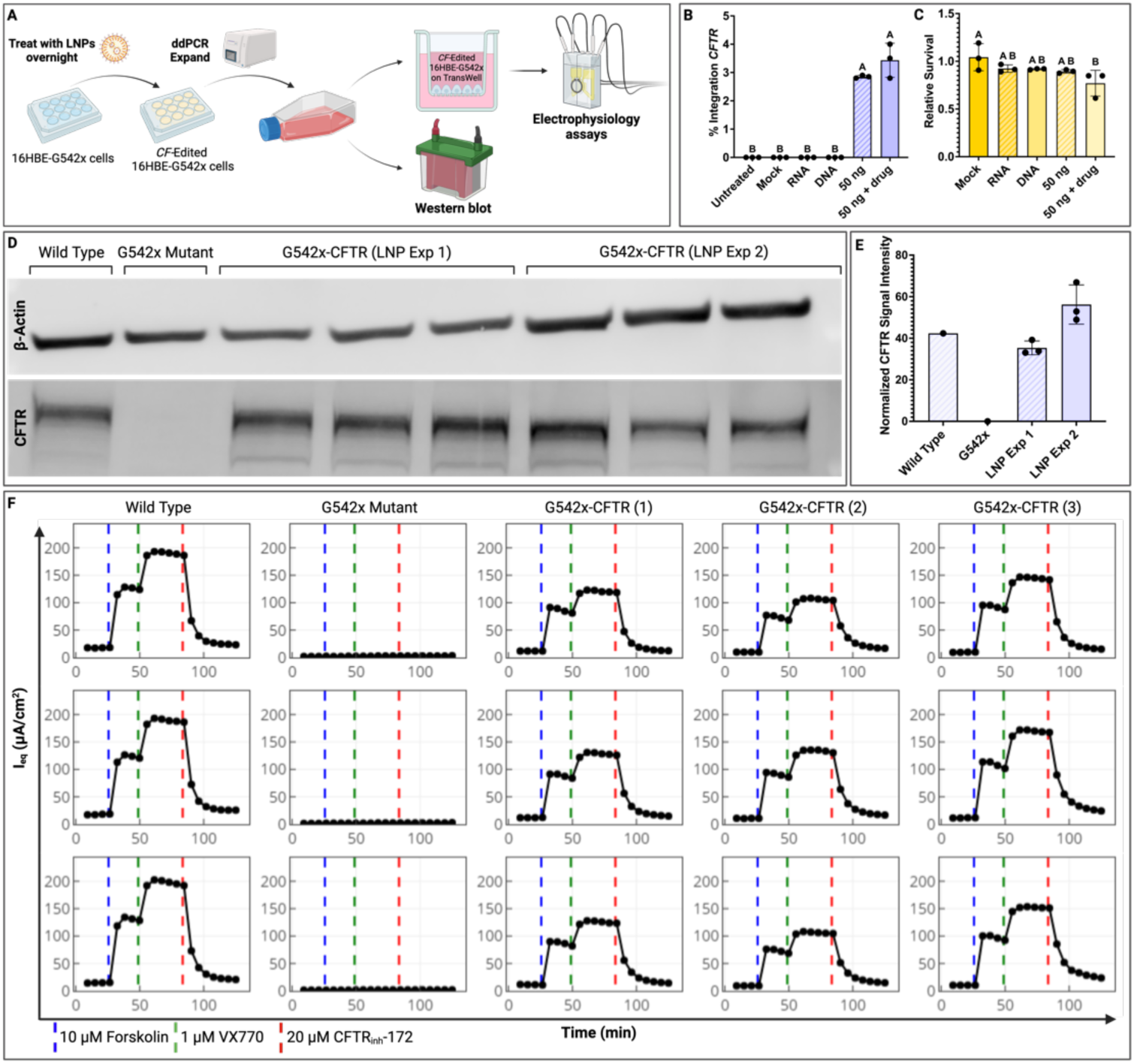
Functional assessment of CFTR-deficient cells edited to express functional CFTR. (**A**) Schematic depicting workflow for LNP-mediated editing of 16HBEgeG542x cells and expansion, and downstream functional analysis of gene edited 16HBEgeG542X cells (G542x-CFTR). (**B**) Droplet digital PCR (ddPCR) data reflecting site-specific integration of a codon-optimized *CFTR* cassette within the 5’UTR of the endogenous *CFTR* gene and (**C**) metabolic tetrazolium salt (MTS) assay data depicting cell survival relative to the untreated condition in G542x-CFTR cells 24 h after treatment with LNPs at various dosages. One-way analyses of variance (ANOVA) were performed, with the threshold of statistical significance set at *p <* 0.05. (**D**) Western blot images depicting protein expression and (**E**) densitometry measurements in cultures expanded from G542x-CFTR cells. (**F**) Transepithelial current clamp (TECC) traces depicting equivalent CFTR-dependent Cl- ion current in CF mutant 16HBEgeG542X cells, wild type 16HBE14o- cells, and G542x-CFTR cells, with each row representing a separate set of technical replicates.

### LNPs encapsulate large dsDNA constructs

Traditional LNPs optimized for RNA delivery have not been fully explored for their ability to encapsulate and deliver larger ldsDNA constructs. Cryogenic transmission electron microscopy (cryo-TEM) was applied to characterize the size and morphology of LNPs containing either Cas9 mRNA and sgRNA only, ldsDNA encoding *CFTR* only, or Cas9 mRNA, sgRNA, and *CFTR-*encoding ldsDNA all together in a single particle (Fig. 8A, Supplementary Figs. S8-S10). All three LNP types primarily contained particles *ca.* 30 – 100 nm with either round or polyhedral morphology and an electron-dense multilamellar core. However, very large (>200 nm) particles with irregular morphology, empty unilamellar liposomes, and large cylindrical and sheet-like structures were also observed in the samples containing ldsDNA (Fig. 8A, Supplementary Figs. S8-S10). This heterogeneity is consistent with broader size distributions and lower encapsulation efficiencies measured in ldsDNA-containing samples *via* DLS and the RiboGreen assay (Fig. 8B,C). Qualitatively, these secondary structures were observed to a greater degree in the samples containing only ldsDNA compared to samples containing both RNA and DNA. Still, these cryo-TEM images verify the formation of multilamellar particles loaded with nucleic acid in all samples, suggesting that large ldsDNA constructs may be packaged within LNPs.

**Figure 8.**
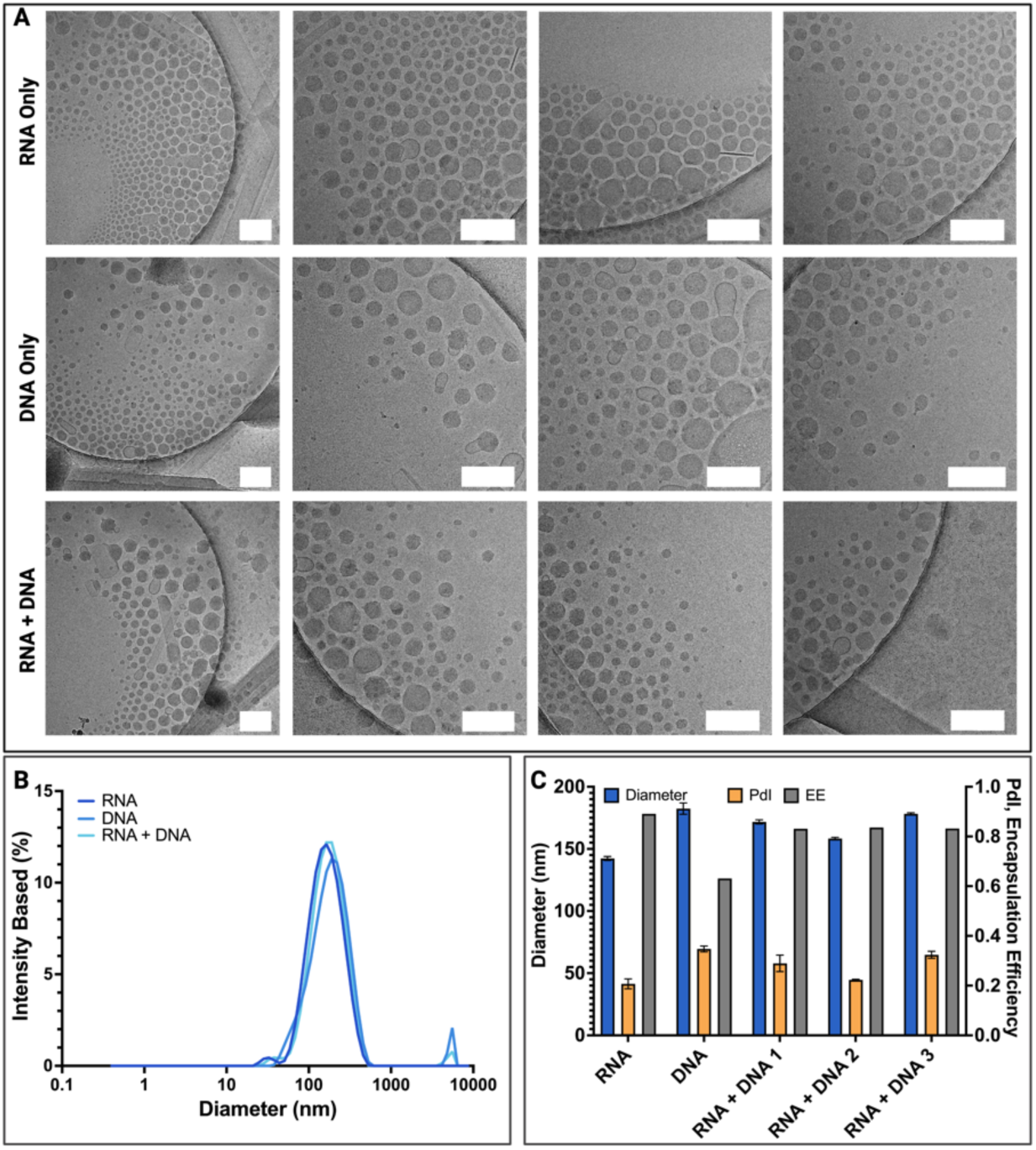
Characterization of lipid nanoparticles (LNPs) loaded with CFTR-editing cargoes. (**A**) Cryogenic transmission electron microscopy (cryo-TEM) images depicting LNPs containing Cas9 mRNA and single guide RNA (sgRNA) (top), linear double-stranded DNA (ldsDNA) encoding CFTR (middle), or Cas9 mRNA, sgRNA, and CFTR ldsDNA (bottom). Scale bars = 200 nm. (**B**) Dynamic light scattering (DLS) plots of the size distribution of LNPs loaded with Cas9 mRNA and sgRNA only, CFTR ldsDNA only, or Cas9 mRNA, sgRNA, and CFTR ldsDNA. (**C**) DLS and RiboGreen encapsulation assay assessment of the Z-ave diameter (blue, left y-axis), polydispersity (PdI) (yellow, right y-axis), and encapsulation efficiency (EE) (grey, right y-axis) of LNPs containing Cas9 mRNA and sgRNA only, CFTR ldsDNA only, or Cas9 mRNA, sgRNA, and CFTR ldsDNA.

## Discussion

Lipid nanoparticles have demonstrated remarkable versatility and widespread clinical success as a platform for siRNA therapeutics and mRNA vaccines, in addition to rapid preclinical development as delivery systems for RNAs of multiple sizes and types, plasmid DNA, and Cas9 RNP complexes (*4–7, 31–33*). However, to date, it remains unclear whether large (>5kb) ldsDNA cassettes are amenable to packaging and intracellular transport by LNPs. In this study, we addressed two key questions: i) can LNPs assembled from commercially available lipid reagents be loaded with and facilitate the delivery of ldsDNA cargoes; and ii) can LNPs be used to coordinate the transport of all components of the CRISPR/Cas9 system to human bronchial epithelial cells to achieve site-specific insertion of HDR donor templates encoding the entire *CFTR* gene and restore protein expression and correct ion channel function in airway epithelia?

To probe the former, we employed DLS, encapsulation assays, and cryo-TEM to compare ldsDNA- containing LNPs with those containing only RNA cargoes. Our data suggest that identical lipid formulations can encapsulate RNA and ldsDNA cargoes alike. However, we observed morphological variations (including the formation of large, irregular LNPs, empty liposomes, and sheet-like structures) in ldsDNA-containing LNPs, suggesting that nanoparticle packaging and assembly are less efficient in these formulations (Fig. 8). We hypothesize that these structural differences may result from altered thermodynamics of self-assembly, perhaps due to greater size and/or rigidity of the ldsDNA donor compared to single-stranded RNA molecules, or because of steric hindrance preventing electrostatic interactions between ionizable lipid amines and the ldsDNA phosphate backbone. Interestingly, samples containing both ldsDNA and RNA tended to exhibit intermediate cargo encapsulation efficiencies and morphological heterogeneity. It remains unclear whether this trend occurs because RNA enhances the self-assembly process to stabilize LNPs containing both RNA and ldsDNA, or if the RNA and ldsDNA partition into separate small, uniform RNA-containing LNPs and larger, heterogenous DNA-containing LNPs. Molecular dynamics studies and advanced analytical techniques, such as the multi-laser cylindrical illumination confocal spectroscopy technique (CICS) may be helpful in elucidating the payload copy number per particle and clarifying these observations (*34*). The morphological heterogeneity of LNPs containing ldsDNA should be further explored with other classes of lipids to determine whether sufficient design and optimization of LNP formulations may enable homogenous ldsDNA encapsulation.

This work demonstrates the feasibility for LNPs to deliver ldsDNA constructs intracellularly for site-specific gene insertion *via* HDR. While *CFTR* integration efficiencies achieved were modest, high (40 – 60%) rates of indels introduced by NHEJ at the cut site in samples edited with both mCitrine and CFTR donor constructs suggest that uptake and endosomal escape are not the limiting factors for our platform (Fig. 5F, Supplementary Materials Fig. S4). Instead, donor trafficking to the nucleus may be inefficient due either to the lack of nuclear localization signal (NLS) or to degradation by cytoplasmic immune machinery. While foreign double-stranded DNA is known to be immunogenic (*35, 36*), cell survival and proliferation observed in this study were not significantly impacted by LNP-mediated delivery of ldsDNA (Fig. 6I, 7C).

Further, the internal biology of the cell likely limits HDR efficiency. This repair process is cell-cycle dependent, occurring primarily in S/G2 phases, and in competition with NHEJ and MMEJ. While this study leverages serum starvation to synchronize cell cycle and applies the recently developed DNA phosphokinase and polymeraseΘ inhibitor drugs AZD-7648 and ART-558 to inhibit NHEJ and MMEJ respectively, these tools are not yet translatable to the clinic (*37, 38*). Broader translation of technologies such as the LNP platform described in this study will require the development of clinically translatable approaches to bias cells toward HDR *in vivo.* Alternatively, methods to increase expression of integrated genes may ameliorate their phenotypic impact, requiring that fewer cells be edited to achieve therapeutic effect. For example, while rates of integration attainable with the HDR-based platform reported here fall below those estimated to achieve phenotypic correction of CF (*39*), data from this study corroborate our recent work demonstrating the use of codon-optimized ldsDNA donor cassettes to enable robust restoration of CFTR ion channel function at integration levels <3% (*23*). While protein translated from codon-optimized transcripts should be carefully assessed for aberrant post-translational modifications and proper protein folding prior to clinical use (*22, 40, 41*), the use of codon-optimized and gain-of-function variants of donor cassettes offer potential solutions for LNP-based systems that must overcome a litany of barriers to efficient intracellular delivery and HDR-based genomic integration. CRISPR-based strategies for genomic integration of entire gene sequences are rapidly advancing toward elegant systems such as CASTs that do not rely on HDR or require DSBs in the endogenous DNA, making them cell cycle independent (*17*). To date, these systems still require ldsDNA donor constructs that could be delivered by LNP platforms such as the one described here, and the synergy of these tools should be explored. Specifically, ionizable lipid libraries and combinatorial LNP formula panels should be further interrogated for their ability to more efficiently encapsulate ldsDNA constructs into uniform particles to advance toward the broader development and standardization of clinically translatable gene therapies.

### Outlook and Future Prospects

This study successfully demonstrates LNP-mediated integration of an entire gene resulting in functional phenotypic correction, establishing a new benchmark for the cargo-carrying capacity of LNPs. These LNPs were applied for the packaging and delivery of ldsDNA donor templates encoding CFTR, which including homology arms totals 5.5 kb in length and is approximately 3400 kDa. LNPs configured with this capability may offer a platform flexible enough to enable delivery of donor templates for the mutation-agnostic correction of any number of single-gene disorders. The adaptation of LNPs as carriers for mRNA began with formulations originally optimized for the delivery of siRNA, and advanced *via* the design and testing of massive libraries of ionizable lipids for encapsulation and endosomal escape profiles that were synergistic with the physicochemical structure of mRNA (*4–7, 42*). A similar multidisciplinary effort may be required to design a class of lipid constituents that are optimized to package ldsDNA constructs for future therapeutic applications. The data presented here offer proof-of-concept that LNPs can mediate the functional delivery of large ldsDNA constructs, indicating that such an effort may be a worthwhile endeavor for researchers developing new LNP-based therapies.

Insight from this work has broad implications for addressing genetic disorders such as CF that arise from any one of thousands of disease-causing mutations. Gene addition strategies, including the CRISPR/Cas9-mediated site-specific insertion approach described in this Article in addition to CAST approaches currently under rapid development for similar applications, offer virtually universal gene correction tools for CF and a litany of other genetic diseases including hemophilia A and adenosine deaminase-deficient severe combined immunodeficiency (ADA-SCID). An existing barrier to clinical translation and implementation of these groundbreaking technologies is the challenge of delivery, and we demonstrate here that LNPs offer an exciting and elegant platform for the intracellular transport of the multiple components required for site-specific whole-gene insertion.

## Materials and Methods

### Chemicals

IL-A, IL-B, IL-C, LP-A, and LP-B were purchased from Echelon Biosciences. Sterol-A, sterol-B, and sterol-C were purchased from Avanti Polar Lipids. LP-C was purchased from NanoSoft Polymers. Chloroform (Sigma) and ethanol (Decon Labs) were purchased from the UCLA Chemistry & Biochemistry Storeroom. Triton^TM^ X-100 was purchased from Millipore Sigma. Quant-iT^TM^ Ribogreen^®^ RNA reagent and ribosomal RNA standards were purchased from Thermo Scientific. CellTiter 96^®^ AQueous One Solution Cell Proliferation Assay was purchased from Promega. EGFP mRNA and Cas9 mRNA were purchased from TriLink Biotechnologies. Single guide RNAs were obtained from Synthego. Phosphate-buffered saline (PBS) was purchased from Thermo Scientific. Citric acid buffer (50 mM, pH 5.35) was prepared by combining 8.55 g sodium citrate dihydrate (Fisher) and 4.02 g citric acid (Fisher) per 1 L diethyl pyrocarbonate (DEPC)-treated water (Thermo Scientific) and adjusting the pH with HCl or NaOH (Sigma-Aldrich). Citric acid buffer (50 mM, pH 4) was made by combining 4.96 g sodium citrate dihydrate and 6.36 g citric acid per 1 L DEPC-treated water and adjusting the pH with HCl or NaOH.

### Nanoparticle formulation, synthesis, and characterization

LNPs containing ionizable lipid (IL-A, IL-B, or IL-C), sterol (sterol-A, sterol-B, or sterol-C), helper lipid (DOPE), and PEG-lipid (LP-A, LP-B, or LP-C) at molar ratios of 50:38.5:10:1.5 respectively were synthesized using a toroidal microfluidic mixing system (NanoAssemblr Spark, Precision NanoSystems). The NanoAssemblr Spark operates at a constant flow rate and 2:1 ratio of aqueous phase to organic phase. A lipid mix concentration of 65 mM was used. LNPs were diluted in PBS. Upon LNP assembly, hydrodynamic size and PdI of particle batches was measured using DLS with a Malvern Zetasizer Nano using a backscatter detection angle of 173°. Dosage of LNPs was determined based on the theoretical concentration of mRNA in the LNP suspension assuming 100% recovery. To determine nucleic acid encapsulation efficiency, LNPs or PBS blanks were diluted in tris-EDTA (TE) buffer to achieve a concentration of 2 – 10 ng/µL nucleic acid per well. These samples were aliquoted and diluted 1:1 in TE buffer (to measure unencapsulated nucleic acid) or TE buffer with 2% Triton-X- 100 (to measure total nucleic acid). Quant-iT RiboGreen reagent was added, and fluorescence signal was quantified with a Varioskan LUX multimode microplate reader (Thermo Scientific). Encapsulation efficiency was calculated as follows:

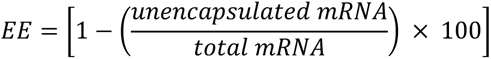

**Equation 1.** Calculation of encapsulation efficiency based on the RiboGreen RNA detection assay.

### Transmission electron microscopy

Samples for cryo-TEM imaging were prepared by depositing 2.5 µL of undiluted LNPs (*ca.* 7 µg/µL lipid or 150 µg/µL nucleic acid) onto a glow-discharged 200 mesh Cu grid with a thin carbon film supported by a holey carbon substrate (QuantiFoil). Grids were blotted for 2 sec at 22°C then plunged into liquid ethane using a manual plunger and transferred to a cryo-TEM grid holder. Imaging was conducted using a K2 camera (Gatan) on a TF20 microscope (FEI) operated at 200 kV.

### Cell Culture

16HBE14o- cells, a human bronchial epithelial cell line with normal CFTR function (*43*), originally developed by Dr. Dieter Gruenert at the University of San Francisco, were obtained from Millipore Sigma. 16HBEge-G542X cells were obtained *via* a materials transfer agreement (MTA) between the Regents of the University of California, Los Angeles and Dr. Hillary Valley at the Cystic Fibrosis Foundation (CFF). All cell lines were cultured in Minimum Essential Medium (MEM) Eagle (Sigma) with 10% fetal bovine serum (FBS) (Millipore Sigma),1% penicillin/streptomycin (Fisher Scientific), and 2 mM L-glutamine (Millipore Sigma) (complete medium referred to as E10). Cells were grown on tissue culture flasks and well plates (Corning) coated with a matrix of bovine serum albumin (BSA, Millipore Sigma), fibronectin from human plasma (Millipore Sigma), and collagen type I (Millipore Sigma) as previously described (*44*). Complete culture medium and coating solution recipes can be found in Supplementary Materials (Tables S1, S2). Cells were passaged by lifting the cells *via* trypsinization with trypsin-EDTA 0.25% (Thermo Fisher Scientific) and split at ratios between 1:4 and 1:10.

### LNP-mediated Transfection

16HBE cells cultured for ten or fewer passages were plated in clear-bottom 96 well plates at 20,000 cells/well. For GFP mRNA delivery experiments, cells were plated in 100 µL E10 and incubated overnight before replacing medium with 100 µL fresh E10 immediately prior to LNP treatment. For gene editing experiments, cells were plated in 100 µL OptiMEM (Thermo Fisher) and incubated for 18 h before replacing medium with 100 µL E10 with or without 0.5 µM AZD-7648 and 2 µM ART-558 (MedChem Express) 4 h prior to adding LNPs. For LNP experiments including *CFTR* donor cassettes, a rho-kinase inhibitor (10 µM Y-27632, STEMCELL Technologies) was added to the cell culture medium 24 h prior to transfection. Cells were cultured with Y-27632 until harvest. Nanoparticles (25 µL) were added directly to the medium and incubated with the cells overnight before replacing medium with 200 µL E10 the next day.

### Flow Cytometry

16HBE14o- and 16HBE14o-BFP were plated in 96-well plates at 20,000 cells/well and treated with LNPs carrying either GFP mRNA or Cas9 mRNA, sgRNA-BFP, and ssODN-BFP. For GFP delivery experiments, cells were harvested the following day *via* trypsinization and resuspended in PBS containing 2% FBS. For BFP to GFP editing experiments, cells were harvested 5 days after transfection *via* trypsinization and resuspended in PBS containing 2% FBS. Cells were stained for viability using 7AAD dye (Thermo Fisher). Analysis was performed on a BD LSRII flow cytometer equipped with 355, 405, 488, 561, and 633 nm lasers and running BD FACS Diva v.8.0.1 software. Forward scatter and side scatter gating was used to exclude debris. Doublets were excluded based on forward scatter width and height gating. GFP was detected through a 525/50 bandpass filter, BFP was detected through a 450/50 bandpass filter, and 7AAD was detected through a 710/50 bandpass filter. Compensation was applied to mitigate spectral overlap between fluorescence signals. Gating was performed as exemplified in Supplementary Materials.

### Live Cell Nuclear Staining and Confocal Microscopy

Complete fixation and immunostaining protocols can be found in the Supplementary Materials. 16HBEs on collagen IV-treated TransWell membranes were incubated in E10 with BioTracker 650 live cell nuclear dye (Millipore Sigma) 18 h prior to imaging. Membranes were rinsed twice with PBS and excised from the supporting inserts with a scalpel. Membranes with live cells carefully sandwiched between a coverslip and glass slide in a droplet of PBS. Cells were imaged using a Leica SP8-STED/FLIM/FCS laser scanning confocal microscope (Leica Microsystems, Wetzlar, Germany).

### DNA Extraction and Analysis

Cells were harvested for DNA extraction 48 – 72 h after transfection *via* trypsinization or cell scraping. Genomic DNA was extracted from cell pellets for all ddPCR and sequencing analyses using the GeneJet Genomic DNA Purification Kit (Thermo Fisher).

### Ussing Chamber Analysis

Cells were cultured and assays were carried out as previously described (*43*). Briefly, normal (16HBE14o-), mutant (G542X), and gene-edited G542X cells from bulk populations (G542X-CFTR) were seeded at a density of 5 x 10^5^ cells on 12 mm polyester SnapWell inserts (#3801; Corning, Corning Inc., NY, USA) pre-coated with human collagen type IV (Sigma-Aldrich, C5533) as described in Supplementary Materials. Cells were grown as submerged cultures in E10, incubated at 37°C and gassed with 5% CO_2_. After a total of 7-9 days, 16HBE cells typically formed electrically tight epithelia. SnapWell inserts were mounted on a slider into water-jacketed Easy Mount Ussing chambers (Physiologic Instruments, Reno, NV). Transepithelial voltage was clamped to 0 mV and resulting short-circuit current (I_sc_) was measured using a four-electrode voltage clamp (VCC MC2 multichannel voltage clamp, Physiologic Instruments), with Ag-AgCl electrodes connected to the solutions through 3% agar bridges (LB Agar; Thermo Fisher) containing 3 M KCl. Short-circuit current was recorded to a computer through an analog-to-digital board (DI710, DataQ Instruments, Akron, OH). At 60 sec intervals, transepithelial voltage was clamped to 1 mV for 1 sec to calculate transepithelial electrical resistance (RT). Current deflections are shown in Ussing traces to visualize RT of each culture. Ussing chamber experiments were performed in HEPES buffered solutions with a serosal-to-mucosal chloride gradient to increase the driving force for chloride exit across the apical cell membrane. The serosal solution contained (mM): 137 NaCl, 4 KCl, 1.8 CaCl_2_, 1 MgCl_2_, 10 HEPES and D-Glucose, adjusted to pH 7.4 with NaOH ([Cl^-^]total: 146.6 mM). The apical solution was matched to the basolateral except for (mM): 137 Na-gluconate replaced 137 NaCl ([Cl^-^]total: 9.6 mM). Full recipes are included in the Supplementary Materials. Fluid volume was 5 ml on each side. Both hemi-chambers were gassed with air and temperature was maintained at 35–37°C. 10 µM amiloride (Selleck Chemicals) was used to block sodium currents. 10 µM forskolin (Tocris) and 1 µM VX-770 (MedChem Express) were applied to activate CFTR chloride currents. CFTR-Inh172 (20-40 µM, MedChem Express) was added for specific block of CFTR-mediated chloride currents. 100 µM adenosine triphosphate (ATP, Thermo Fisher) was used to stimulate calcium-activated Cl currents, which served as a quality control for each culture. Agonists/antagonists were added to both the basolateral and apical sides of the Ussing reservoir. Changes in I_sc_ in response to CFTRinh172 were used as a measure of functional CFTR surface expression and treatment-related functional correction. Normal, 100% CFTR activity was defined as maximal *Δ*I_sc_ in response to CFTRinh172 in normal 16HBE14o- cells.

### Transepithelial Current Clamp Assays

The protocol used for TECC studies was adapted from Valley and colleagues (*44*). 16HBE cells were seeded at a density of 4.5 × 10^5^ cells/cm^2^ onto HTS TransWell 24-well filter inserts (Corning, 3378) pre-coated with human collagen type IV (Sigma-Aldrich, C5533). Cells were grown as submerged cultures in MEM (Gibco, 11095) containing 10% FBS (Hyclone, SH30071.03) and 1% Pen/Strep, and incubated at 37°C and 5% CO_2_. After a total of 7 days, 16HBE cells typically formed electrically tight epithelia with a transepithelial resistance (Rt) of <800 Ω·cm^2^ and CFTR-mediated Cl^-^ equivalent current (I_eq_) was determined as described below. Prior to functional (I_eq_) studies, MEM was replaced with fresh HEPES-buffered (pH 7.4) solutions (assay buffer). A driving force for chloride ions was established through application of a basolateral to apical chloride ion gradient (see buffer composition below). Cell plates were mounted onto an automated robotic assay platform and equilibrated at ∼36°C for 90 min. After equilibration, transepithelial voltage (Vt) and resistance (Rt) were monitored at ∼5 min intervals using a 24-channel transepithelial current clamp amplifier (TECC-24, EP Design, Bertem, Belgium). Electrode potential differences for each pair of Ag/AgCl voltage electrodes were also monitored at 5 min intervals by taking voltage measurements from a control plate with matching buffer solutions and 16HBE cells that were left untreated. I_eq_ was calculated from values of Vt and Rt using Ohm’s law after correcting for series resistance and (electrode) voltage offsets unrelated to Vt. I_eq_ traces are plotted as mean ± SD (n = 3). The first 4 data points reflect baseline I_eq_ currents prior to sequential stimulation of CFTR with forskolin (10 μM) and VX-770/ivacaftor (1 μM). The last six data points were recorded in the presence of CFTR inhibitor CFTRinh-172 (20 μM). Agonists/antagonists were pre-diluted to 10-fold concentrations in assay buffer and added to either the basolateral (forskolin) or apical (forskolin, VX-770/ ivacaftor, and CFTRinh-172) side of the membrane (assay plates only). CFTR-mediated changes in I_eq_, (*i.e.,* delta forskolin, delta VX-770, delta CFTRinh-172, or the area under the curve (AUC) between forskolin and CFTRinh-172 addition) are used as a measure of functional CFTR surface expression or treatment-related functional rescue of mutant CFTR.

CFTR-mediated transepithelial currents were recorded using a Cl^-^ concentration gradient. The basolateral solution contained (mM): 137 NaCl, 4 KCl, 1.8 CaCl2, 1 MgCl2, 10 HEPES and D-Glucose, adjusted to pH 7.4 with NaOH/HCl ([Cl^-^]total: 146.6 mM). The apical solution was matched to the basolateral except for (mM): 137 Na-gluconate replaced 137 NaCl ([Cl^-^]total: 9.6 mM). Full buffer recipes are included in the Supplementary Materials.

### Western Blot

For immunoblots, cells were lysed in RIPA Lysis and Extraction Buffer (Cat: 89901; ThermoFisher Scientific; Grand Island, NY) with added HALT protease inhibitor (Cat: 87786; ThermoFisher Scientific; Grand Island, NY) at a 1 × concentration following the manufacturer’s protocols. Lysate concentrations were determined using the Pierce BCA protein assay (Cat: 23227; ThermoFisher Scientific, Grand Island, NY) following the manufacturer’s protocol. Samples were treated for sodium dodecyl sulfate–polyacrylamide gel electrophoresis (SDS-PAGE) with NuPAGE LDS Sample Buffer (Cat: NP0007; ThermoFisher Scientific; Grand Island, NY) and NuPAGE Sample Reducing Agent (Cat: NP0009; ThermoFisher Scientific; Grand Island, NY), each to a 1 × concentration. Lysates were diluted to contain 50 μg of total protein for immunoblot gel loading to keep the total amount of protein loaded per lane constant to allow for valid loading controls. Note, CFTR protein becomes insoluble if the sample is heated above 60°C and will not enter the stacking gel. As such, the samples were denatured at 37°C for 20 min prior to loading into the stacking gel. Wild-type cells 16HBE14o- were used as a control to indicate the relative expression levels of CFTR protein. CFTR levels were detected using Ab 596 (obtained *via* MTA from J. Riordan, UNC;) at 1:1000 in 5% milk in TBST (*45*). Protein quantification was assessed through densitometry *via* the ImageJ software. CFTR protein levels were normalized to the actin protein levels after quantification.

### Double stranded DNA donor synthesis and amplification

Linear double-stranded DNA (ldsDNA) donor fragments were synthesized as gBlocks by Integrated DNA Technologies (Coralville, IA). The gBlocks were subsequently cloned into plasmids using the TOPO Zero Blunt Cloning Kit (Thermo Fisher Scientific, Catalog #K2800J10). The ldsDNA donors were then amplified by PCR from these plasmids with Platinum SuperFi II DNA Polymerase using the manufacturer’s protocol (ThermoFisher; Catalog #12369010). Amplification utilized oligonucleotide primers “GTCTTTGGCATTAGGAGCTT” and “AGACAACGCTGGCCTTTTC” that were modified at the 5’ end with AmC6, also supplied by Integrated DNA Technologies (Coralville, IA). The PCR products were then purified using SPRI paramagnetic bead-based purification (complete protocol and ldsDNA sequences are provided in Supplementary Materials).

### Integration analysis by ddPCR

Genomic DNA was extracted from edited cells for integration site analysis using the Invitrogen PureLink Genomic DNA Kit (Cat: K182002; Thermo fisher Scientific) and quantified using a NanoDrop system (Cat: ND- 2000; Thermo Fisher Scientific). DNA samples were then analyzed by ddPCR to measure integration rates. Two sets of primers were duplexed, each with their own fluorescent probe (FAM/HEX, Table 1). To measure integration rates of the mCitrine reporter cassette, one primer was complementary to a CFTR gene sequence upstream from the left homology arm of the donor. The second primer bound to the mCitrine reporter cassette, to allow for specific measurement of the integrated mCitrine transgene distinct from the endogenous CFTR gene. To measure integration rates of the CFTR donor cassette, one primer was bound to the bovine growth hormone polyA signal sequence to allow for specific measurement of the integrated CFTR donor. The second primer was complementary to a CFTR gene sequence upstream from the right homology arm of the donor cassette. The FAM-conjugated nucleotide probe with a quencher also bound to the minus-strand DNA near the second primer. A reference primer/probe set was also delivered to recognize the SDC4 gene on chromosome 20. One microliter of DraI endonuclease (Cat: R0129S; New England Biolabs) was added to the reaction mixture (an enzyme that does not disrupt the experimental or reference amplicons) to reduce background. Each sample was digested at 37°C for 1 h before droplet generation with the Bio-Rad QX200 Droplet Generator (Cat: 186-4002; Bio-Rad, Hercules, CA). The droplet emulsion was then transferred with a multichannel pipet to a 96-well twin.tec® real-time PCR Plate (Eppendorf; Hamburg, Germany) heat sealed with foil, and amplified in a conventional thermal cycler (T100 Thermal Cycler; Bio-Rad; Hercules, CA). Thermal cycling conditions consisted of 96°C 10 min (1 cycle), 94°C 30 s and 60°C 1 min (55 cycles), 98°C 10 min (1 cycle) and 12°C hold. After PCR, the 96-well plate was transferred to the QX200 Bio-Rad Droplet Reader on the “Absolute” measurement setting (Cat: 186-4003; Bio-Rad). Acquisition and analysis of the ddPCR data was performed with the QuantaSoft software (Bio-Rad; Hercules, CA).

**Table 1.**
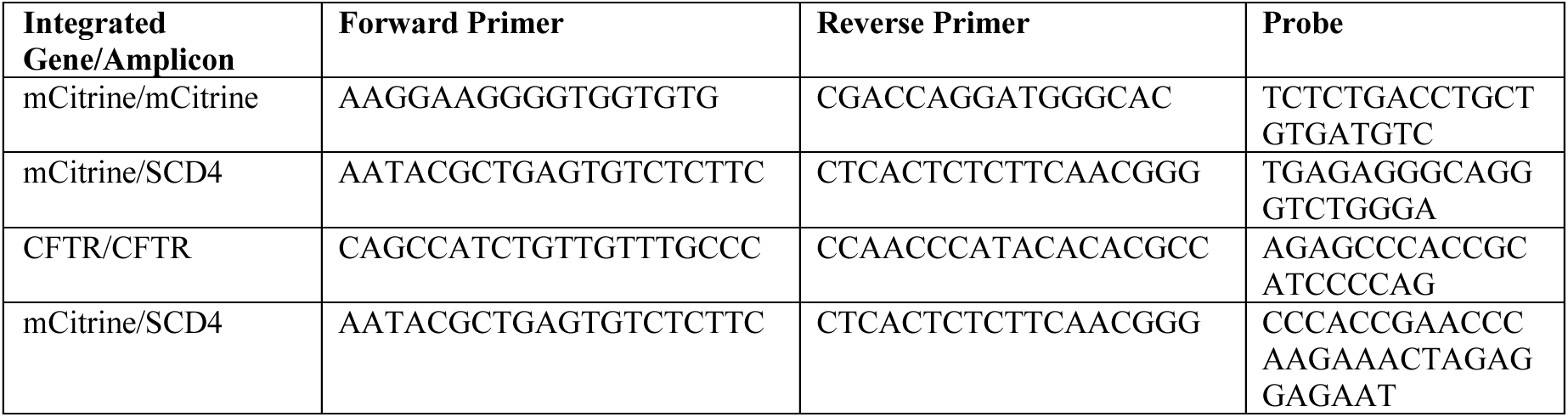
Primer/probe sets used for the detection of integrated sequences using Droplet digital PCR (ddPCR).

### Measuring allelic disruption

To measure allelic disruption of the reporter gene cassette, a 728 bp amplicon that encompasses CFTR 5’UTR and exon 1 was amplified from genomic DNA taken from edited cells (primer sequences: “AAAGCCGCTAGAGCAAATTT,” “TGTTGGCTGAATTCAGTCAA”). The resulting amplicon was Sanger sequenced and uploaded for analysis *via* the Synthego ICE web tool (Synthego Corporation, Menlo Park, CA).

### Generation of BFP to GFP 14HBE16o- Cell Line

16HBE14o- cells were transduced with a lentivirus carrying a BFP reporter cassette driven by the EF1*α*promoter based on a protocol described by Dr. Corn and colleagues (Addgene plasmid #71825; https://www.addgene.org/71825/) (*26*). Following transduction, BFP-expressing cells were identified and quantified using flow cytometry. Single-cell sorting of BFP-positive clones was performed using the BD FACS ARIA sorter (BD Biosciences; Franklin Lakes, NJ). The sorted clones were expanded in culture, and their vector copy number was determined using ddPCR. This was achieved by quantifying the HIV-1 PSI region of the lentiviral vector relative to the endogenous human diploid control gene, SDC4, as previously described (*46*). Clones with a vector copy number of 1 were selected for further experiments and designated as the BFP 16HBE14o- cell line.

### Statistical analysis

Statistical analyses were performed using GraphPad Prism 9 software. Data sets were checked for normality using a Shapiro-Wilk test. Population comparisons from data sets passing normality tests (alpha = 0.05) were analyzed using a one-way analysis of variance (ANOVA). Population comparisons from data sets not passing normality tests were analyzed using a Kruskal-Wallis test of variance. The threshold for statistical significance was set at *p =* 0.05. All error bars denote standard deviation.

## Acknowledgments

The authors acknowledge the use of instruments and core resources made available at the Electron Imaging Center for Nanosystems (EICN) supported by NIH (1S10RR23057, 1S10OD018111 and 1U24GM116792), NSF (DBI-1338135) and California NanoSystems Institute (CNSI) at UCLA. We further acknowledge the use of instruments at the Nano and Pico Characterization Lab at CNSI. Confocal laser scanning microscopy was performed at the CNSI Advanced Light Microscopy/Spectroscopy (ALMS) Laboratory and Leica Microsystems Center of Excellence at the California NanoSystems Institute at UCLA (RRID:SCR_022789) with funding support from NIH Shared Instrumentation Grant S10OD025017 and NSF Major Research Instrumentation grant CHE-0722519. We thank the staff of the BSCRC Flow Cytometry and Microscopy Cores and the EICN, ALMS, and NPC core facilities at CNSI for their valuable help and support, and in particular Dr. Wong Hoi Hui who provided assistance performing cryo-TEM.

## Author contributions

Conceptualization: RAF, PGA, VS, CJJ, BNG, DBK, SJJ

Methodology: RAF, PGA, VS, CJJ, AS, ECD, LEL, PB, KC, BI, BNG, DBK, SJJ

Investigation: RAF, PGA, VS, CJJ, AS, ECD, LEL, PB, KC, BI

Visualization: RAF, PB, KC

Supervision: BI, BNG, DBK, SJJJ

Writing—original draft: RAF

Writing—review & editing: RAF, PGA, VS, CJJ, AS, ECD, LEL, PB, KC, BI, BNG, DBK, SJJ

## Data and materials availability

The 16HBEgeG542X cell line was obtained *via* an MTA (H. Valley, CFF). Similarly, the CFTR antibody Ab 596 used for Western blots were obtained *via* an MTA (J. Riordan, UNC). All data are available in the main text or the Supplementary Materials.

## Supplementary Materials

Supplementary Methods

Tables S1 to S6

Figs. S1 to S10

References

## References

1. D. B. Kohn, C. Y. Kuo, New frontiers in the therapy of primary immunodeficiency: From gene addition to gene editing. Journal of Allergy and Clinical Immunology 139, 726–732 (2017).

2. M. C. Canver, S. H. Orkin, Customizing the genome as therapy for the β-hemoglobinopathies. Blood 127, 2536–2545 (2016).

3. J. A. Doudna, E. Charpentier, The new frontier of genome engineering with CRISPR-Cas9. Science 346, 1258096 (2014).

4. R. A. Foley, R. A. Sims, E. C. Duggan, J. K. Olmedo, R. Ma, S. J. Jonas, Delivering the CRISPR/Cas9 system for engineering gene therapies: Recent cargo and delivery approaches for clinical translation. Frontiers in Bioengineering and Biotechnology 10, 973326 (2022).

5. A. Sengupta, in Viral Infections and Antiviral Therapies. (Elsevier, 2023), pp. 611–624.

6. Y. Suzuki, H. Ishihara, Difference in the lipid nanoparticle technology employed in three approved siRNA (Patisiran) and mRNA (COVID-19 vaccine) drugs. Drug Metabolism and Pharmacokinetics 41, 100424 (2021).

7. J. A. Kulkarni, D. Witzigmann, S. Chen, P. R. Cullis, R. van der Meel, Lipid Nanoparticle Technology for Clinical Translation of siRNA Therapeutics. Accounts of Chemical Research 52, 2435–2444 (2019).

8. A. L. Cooney, P. B. McCray, Jr., P. L. Sinn, Cystic Fibrosis Gene Therapy: Looking Back, Looking Forward. Genes (Basel) 9, 538 (2018).

9. D. Goetz, C. L. Ren, Review of Cystic Fibrosis. Pediatr Ann 48, e154–e161 (2019).

10. A. Berical, R. E. Lee, S. H. Randell, F. Hawkins, Challenges Facing Airway Epithelial Cell-Based Therapy for Cystic Fibrosis. Frontiers in Pharmacology 10, 74 (2019).

11. G. A. Duncan, J. Jung, J. Hanes, J. S. Suk, The Mucus Barrier to Inhaled Gene Therapy. Mol Ther 24, 2043–2053 (2016).

12. J. V. Fahy, B. F. Dickey, Airway mucus function and dysfunction. N Engl J Med 363, 2233–2247 (2010).

13. M. E. McGarry, S. A. McColley, Cystic fibrosis patients of minority race and ethnicity less likely eligible for CFTR modulators based on CFTR genotype. Pediatric pulmonology. 56, 1496–1503 (2021).

14. T. Wei, Y. Sun, Q. Cheng, S. Chatterjee, Z. Traylor, L. T. Johnson, M. L. Coquelin, J. Wang, M. J. Torres, X. Lian, X. Wang, Y. Xiao, C. A. Hodges, D. J. Siegwart, Lung SORT LNPs enable precise homology-directed repair mediated CRISPR/Cas genome correction in cystic fibrosis models. Nature Communications 14, 7322 (2023).

15. Y. Sun, S. Chatterjee, X. Lian, Z. Traylor, S. R. Sattiraju, Y. Xiao, S. A. Dilliard, Y.-C. Sung, M. Kim, S. M. Lee, S. Moore, X. Wang, D. Zhang, S. Wu, P. Basak, J. Wang, J. Liu, R. J. Mann, D. F. LePage, W. Jiang, S. Abid, M. Hennig, A. Martinez, B. A. Wustman, D. J. Lockhart, R. Jain, R. A. Conlon, M. L. Drumm, C. A. Hodges, D. J. Siegwart, In vivo editing of lung stem cells for durable gene correction in mice. Science 384, 1196–1202 (2024).

16. X. Wang, G. Xu, W. A. Johnson, Y. Qu, D. Yin, N. Ramkissoon, H. Xiang, L. Cong, Long sequence insertion via CRISPR/Cas gene-editing with transposase, recombinase, and integrase. Current Opinion in Biomedical Engineering 28, 100491 (2023).

17. G. D. Lampe, R. T. King, T. S. Halpin-Healy, S. E. Klompe, M. I. Hogan, P. L. H. Vo, S. Tang, A. Chavez, S. H. Sternberg, Targeted DNA integration in human cells without double-strand breaks using CRISPR-associated transposases. Nature Biotechnology 42, 87–98 (2024).

18. S. Inouye, Y. Sahara-Miura, J.-i. Sato, T. Suzuki, Codon optimization of genes for efficient protein expression in mammalian cells by selection of only preferred human codons. Protein Expression and Purification 109, 47–54 (2015).

19. H. Bihler, A. Sivachenko, L. Millen, P. Bhatt, A. T. Patel, J. Chin, V. Bailey, I. Musisi, A. LaPan, N. E. Allaire, J. Conte, N. R. Simon, A. S. Magaret, K. S. Raraigh, G. R. Cutting, W. R. Skach, R. J. Bridges, P. J. Thomas, M. Mense, In vitro modulator responsiveness of 655 CFTR variants found in people with cystic fibrosis. Journal of Cystic Fibrosis 23, 664–675 (2024).

20. M. Woodall, R. Tarran, R. Lee, H. Anfishi, S. Prins, J. Counsell, P. Vergani, S. Hart, D. Baines, Expression of gain-of-function CFTR in cystic fibrosis airway cells restores epithelial function better than wild-type or codon-optimized CFTR. Molecular Therapy-Methods & Clinical Development 30, 593–605 (2023).

21. S. Somanathan, F. Jacobs, Q. Wang, A. L. Hanlon, J. M. Wilson, D. J. Rader, AAV Vectors Expressing LDLR Gain-of-Function Variants Demonstrate Increased Efficacy in Mouse Models of Familial Hypercholesterolemia. Circulation Research 115, 591–599 (2014).

22. A. I. Paremskaia, A. A. Kogan, A. Murashkina, D. A. Naumova, A. Satish, I. S. Abramov, S. G. Feoktistova, O. N. Mityaeva, A. A. Deviatkin, P. Y. Volchkov, Codon-optimization in gene therapy: promises, prospects and challenges. Frontiers in Bioengineering and Biotechnology 12, 1371596 (2024).

23. P. G. Ayoub, V. Sinha, C. Juett, L. Lathrop, J. D. Long, R. A. Foley, E. C. Duggan, B. Illek, B. N. Gomperts, S. J. Jonas, D. B. Kohn. (In Preparation).

24. H. Yu, A. Angelova, B. Angelov, B. Dyett, L. Matthews, Y. Zhang, M. El Mohamad, X. Cai, S. Valimehr, C. J. Drummond, J. Zhai, Real-Time pH-Dependent Self-Assembly of Ionisable Lipids from COVID-19 Vaccines and In Situ Nucleic Acid Complexation. Angewandte Chemie. 62, (2023).

25. E. J. Sayers, S. E. Peel, A. Schantz, R. M. England, M. Beano, S. M. Bates, A. S. Desai, S. Puri, M. B. Ashford, A. T. Jones, Endocytic Profiling of Cancer Cell Models Reveals Critical Factors Influencing LNP-Mediated mRNA Delivery and Protein Expression. Molecular Therapy 27, 1950–1962 (2019).

26. C. D. Richardson, G. J. Ray, M. A. DeWitt, G. L. Curie, J. E. Corn, Enhancing homology-directed genome editing by catalytically active and inactive CRISPR-Cas9 using asymmetric donor DNA. Nature biotechnology 34, 339–344 (2016).

27. F.-M. Cloarec-Ung, J. Beaulieu, A. Suthananthan, B. Lehnertz, G. Sauvageau, H. M. Sheppard, D. J. H. F. Knapp, Near-perfect precise on-target editing of human hematopoietic stem and progenitor cells. eLife 12, RP91288 (2024).

28. J. Schimmel, N. Muñoz-Subirana, H. Kool, R. van Schendel, S. van der Vlies, J. A. Kamp, F. M. S. de Vrij, S. A. Kushner, G. C. M. Smith, S. J. Boulton, M. Tijsterman, Modulating mutational outcomes and improving precise gene editing at CRISPR-Cas9-induced breaks by chemical inhibition of end-joining pathways. Cell Reports 42, 112019 (2023).

29. S. Selvaraj, W. N. Feist, S. Viel, S. Vaidyanathan, A. M. Dudek, M. Gastou, S. J. Rockwood, F. K. Ekman, A. R. Oseghale, L. Xu, M. Pavel-Dinu, S. E. Luna, M. K. Cromer, R. Sayana, N. Gomez-Ospina, M. H. Porteus, High-efficiency transgene integration by homology-directed repair in human primary cells using DNA-PKcs inhibition. Nature Biotechnology 42, 731–744 (2024).

30. Cystic Fibrosis Foundation Patient Registry 2023 Annual Data Report. Available from: https://www.cff.org/medical-professionals/patient-registry

31. S. H. Im, M. Jang, J.-H. Park, H. J. Chung, Finely tuned ionizable lipid nanoparticles for CRISPR/Cas9 ribonucleoprotein delivery and gene editing. Journal of Nanobiotechnology 22, 175 (2024).

32. T. Wei, Q. Cheng, Y.-L. Min, E. N. Olson, D. J. Siegwart, Systemic nanoparticle delivery of CRISPR-Cas9 ribonucleoproteins for effective tissue specific genome editing. Nature Communications 11, 3232 (2020).

33. P. H. D. M. Prazeres, H. Ferreira, P. A. C. Costa, W. da Silva, M. T. Alves, M. Padilla, A. Thatte, A. K. Santos, A. O. Lobo, A. Sabino, H. L. Del Puerto, M. J. Mitchell, P. P. G. Guimaraes, Delivery of Plasmid DNA by Ionizable Lipid Nanoparticles to Induce CAR Expression in T Cells. International Journal of Nanomedicine 18, 5891–5904 (2023).

34. S. Li, Y. Hu, A. Li, J. Lin, K. Hsieh, Z. Schneiderman, P. Zhang, Y. Zhu, C. Qiu, E. Kokkoli, T.-H. Wang, H.-Q. Mao, Payload distribution and capacity of mRNA lipid nanoparticles. Nature Communications 13, 5561 (2022).

35. V. Hornung, E. Latz, Intracellular DNA recognition. Nature Reviews Immunology 10, 123–130 (2010).

36. B. R. Shy, V. S. Vykunta, A. Ha, A. Talbot, T. L. Roth, D. N. Nguyen, W. G. Pfeifer, Y. Y. Chen, F. Blaeschke, E. Shifrut, S. Vedova, M. R. Mamedov, J.-Y. J. Chung, H. Li, R. Yu, D. Wu, J. Wolf, T. G. Martin, C. E. Castro, L. Ye, J. H. Esensten, J. Eyquem, A. Marson, High-yield genome engineering in primary cells using a hybrid ssDNA repair template and small-molecule cocktails. Nature Biotechnology 41, 521–531 (2023).

37. B. S. Lapa, M. I. Costa, D. Figueiredo, J. Jorge, R. Alves, A. R. Monteiro, B. Serambeque, M. Laranjo, M. F. Botelho, I. M. Carreira, A. B. Sarmento-Ribeiro, A. C. Gonçalves, AZD-7648, a DNA-PK Inhibitor, Induces DNA Damage, Apoptosis, and Cell Cycle Arrest in Chronic and Acute Myeloid Leukemia Cells. International Journal of Molecular Sciences 24, 15331 (2023).

38. D. Zatreanu, H. M. R. Robinson, O. Alkhatib, M. Boursier, H. Finch, L. Geo, D. Grande, V. Grinkevich, R. A. Heald, S. Langdon, J. Majithiya, C. McWhirter, N. M. B. Martin, S. Moore, J. Neves, E. Rajendra, M. Ranzani, T. Schaedler, M. Stockley, K. Wiggins, R. Brough, S. Sridhar, A. Gulati, N. Shao, L. M. Badder, D. Novo, E. G. Knight, R. Marlow, S. Haider, E. Callen, G. Hewitt, J. Schimmel, R. Prevo, C. Alli, A. Ferdinand, C. Bell, P. Blencowe, C. Bot, M. Calder, M. Charles, J. Curry, T. Ekwuru, K. Ewings, W. Krajewski, E. MacDonald, H. McCarron, L. Pang, C. Pedder, L. Rigoreau, M. Swarbrick, E. Wheatley, S. Willis, A. C. Wong, A. Nussenzweig, M. Tijsterman, A. Tutt, S. J. Boulton, G. S. Higgins, S. J. Pettitt, G. C. M. Smith, C. J. Lord, Polθ inhibitors elicit BRCA-gene synthetic lethality and target PARP inhibitor resistance. Nature Communications 12, 3636 (2021).

39. L. Zhang, B. Button, S. E. Gabriel, S. Burkett, Y. Yan, M. H. Skiadopoulos, Y. L. Dang, L. N. Vogel, T. McKay, A. Mengos, R. C. Boucher, P. L. Collins, R. J. Pickles, CFTR Delivery to 25% of Surface Epithelial Cells Restores Normal Rates of Mucus Transport to Human Cystic Fibrosis Airway Epithelium. PLOS Biology 7, e1000155 (2009).

40. C.-H. Yu, Y. Dang, Z. Zhou, C. Wu, F. Zhao, Matthew S. Sachs, Y. Liu, Codon Usage Influences the Local Rate of Translation Elongation to Regulate Co-translational Protein Folding. Molecular Cell 59, 744–754 (2015).

41. V. P. Mauro, S. A. Chappell, A critical analysis of codon optimization in human therapeutics. Trends in molecular medicine 20, 604–613 (2014).

42. F. Freitag, E. Wagner, Optimizing synthetic nucleic acid and protein nanocarriers: The chemical evolution approach. Advanced Drug Delivery Reviews 168, 30–54 (2021).

43. B. Illek, R. Maurisse, L. Wahler, K. Kunzelmann, H. Fischer, D. C. Gruenert, Cl transport in complemented CF bronchial epithelial cells correlates with CFTR mRNA expression levels. Cell Physiol Biochem 22, 57–68 (2008).

44. H. C. Valley, K. M. Bukis, A. Bell, Y. Cheng, E. Wong, N. J. Jordan, N. E. Allaire, A. Sivachenko, F. Liang, H. Bihler, P. J. Thomas, J. Mahiou, M. Mense, Isogenic cell models of cystic fibrosis-causing variants in natively expressing pulmonary epithelial cells. J Cyst Fibros 18, 476–483 (2019).

45. L. Cui, L. Aleksandrov, X.-B. Chang, Y.-X. Hou, L. He, T. Hegedus, M. Gentzsch, A. Aleksandrov, W. E. Balch, J. R. Riordan, Domain Interdependence in the Biosynthetic Assembly of CFTR. Journal of Molecular Biology 365, 981–994 (2007).

46. P. G. Ayoub, J. Gensheimer, L. Lathrop, C. Juett, J. Quintos, K. Tam, J. Reid, F. Ma, C. Tam, G. E. McAuley, D. Brown, X. Wu, R. Zhang, K. Bradford, R. P. Hollis, G. M. Crooks, D. B. Kohn, Lentiviral Vectors for Precise Expression to Treat X-Linked Lymphoproliferative Disease. Molecular Therapy Methods & Clinical Development 32, 101323 (2024).

