## Supplementary Materials for "Lipid Nanoparticles for the Delivery of CRISPR/Cas9 Machinery to Enable Site-Specific Integration of CFTR and Mutation-Agnostic Disease Rescue"

**This PDF file includes:**

Supplementary Methods
Tables S1 to S5
Figs. S1 to S10
References

### Supplementary Methods

#### Cell Culture

**Table S1.** 16HBE14o- and 16HBEgeG542X cell culture medium components. Complete medium was stored at 4ºC (*1*).

| Component | Vendor/Catalog | % (final) |
|---|---|---|
| Minimum Essential Medium | Millipore Sigma M2279 | 89% |
| Fetal bovine serum | Thermo Fisher 16140089 | 10% |
| Penicillin/Streptomycin (100x) | Thermo Fisher 15-140-122 | 1% |

**Table S2.** Flask and well plate coating solution for culturing 16HBE14o- and 16HBEgeG542X cells (*1*).

| Component | Vendor/Catalog | 50 mL final solution |
|---|---|---|
| Minimum Essential Medium | Millipore Sigma M2279 | 48 mL |
| Bovine serum albumin 7.5% | Millipore Sigma 126575 | 67 µL |
| Bovine collagen solution, Type 1 | Advanced BioMatrix 5005-100ML | 0.5 mL |
| Fibronectin from human plasma, 1mg/ml | Sigma-Aldrich F2006 | 0.5 mL |

*Cultureware coating protocols*

For standard culture, tissue culture surfaces were wetted by the solution described in Table S2. (5 mL for a T75, 1 mL for a T25, 50 µL for a 96 well plate) and left for 2-3 h at 37ºC. After incubation, coating solution was aspirated and cultureware was used immediately or stored at 4ºC.

For electrophysiology studies, 12 mm polyester SnapWell inserts (Corning) were wetted with 200 µL collagen type IV solution from human placenta (Millipore Sigma, catalog #C7521). Stock solutions of collagen were prepared by adding 10 mL tissue culture-grade double-distilled water ($ddH_20$) to 5 mg collagen, then adding 20 µL glacial acetic acid. The solution was incubated at 37ºC for 15-30 min until dissolved. This stock solution was diluted 1:10 in ddH20 prior to coating SnapWell inserts. Inserts were incubated at room temperature overnight. Any remaining liquid was aspirated, and the inserts were UV sterilized for 30 min prior to use.

**Using Chamber Buffer Recipes**

**For gradient assay:**

**Table S3.** Equilibration buffer recipe (*1*).

| Solution (1M) | Amt (mL) |
|---|---|
| NaCl | 137 |
| KCl | 4 |
| $CaCl_2$ | 1.8 |
| $MgCl_2$ | 1 |
| HEPES | 10 |
| Adjust volume to 1 L with ddH2O and pH to 7.4 with 1M NaOH. Sterile filter. | |

**Table S4.** Serosal buffer recipe (High $Cl^-$) (*1*).

| Solution (1M) | Amt (mL) |
|---|---|
| NaCl | 137 |
| KCl | 4 |
| $CaCl_2$ | 1.8 |
| $MgCl_2$ | 1 |
| HEPES | 10 |
| D-Glucose | 10 |
| Adjust volume to 1 L with ddH2O and pH to 7.4 with 1M NaOH. Sterile filter. | |

**Table S5.** Mucosal buffer recipe (Low $Cl^-$) (*1*).

| Solution | Amt (mL) |
|---|---|
| Na-gluconate | 137 |
| KCl | 4 |
| CaCl2 | 1.8 |
| MgCl2 | 1 |
| HEPES | 10 |
| D-Glucose | 10 |
| Adjust volume to 1 L with ddH2O and pH to 7.4 with 1M N-methyl-D-glucamine (NMDG). Sterile filter. | |

**Double stranded DNA Bead Purification**

To begin the PCR purification process, the PCR reaction mixture was divided into 500 µL portions. Beads were vortexed, then an equal volume of 1.0x SPRI paramagnetic beads (Ampure XP; Beckman Coulter; # A63881) were added to the PCR reaction. The suspension was mixed thoroughly by pipetting up and down ten times. The mixture was allowed to sit for 5 min before being placed on a magnet until the liquid clarified. The supernatant was carefully removed without disturbing the bead pellet. With the tube on the magnetic plate, the beads were washed with at least 1 mL of 70% ethanol, ensuring the beads are covered but the pellet remained undisturbed on the magnet. The ethanol was allowed to incubate for at least 10-30 sec at room temperature before removing the supernatant. This washing step was repeated with another 1 mL of 70% ethanol. After removing all ethanol, the beads were rehydrated by adding 150 µL of 1/10th Tris-EDTA (TE) buffer, mixing thoroughly with a pipette, and then incubating for 5 min. The tube was returned to the magnet, and the supernatant was extracted once the solution clarified. Another 150 µL of 1/10th TE was added, and the suspension was mixed, incubated, and then the supernatant was extracted to combine with the previous extract. The DNA concentration was quantified using a NanoDrop (Thermo Fisher). Next, 30 µL of CutSmart and 7 µL of Dpn1 were added to the tube, incubating at 37°C for 1 h. For a second round of purification, 340 µL of SPRI paramagnetic beads were used without dividing the sample. The DNA was eluted in 80 µL of pure TE, and the concentration was quantified using a NanoDrop.

**Supplementary Figures**

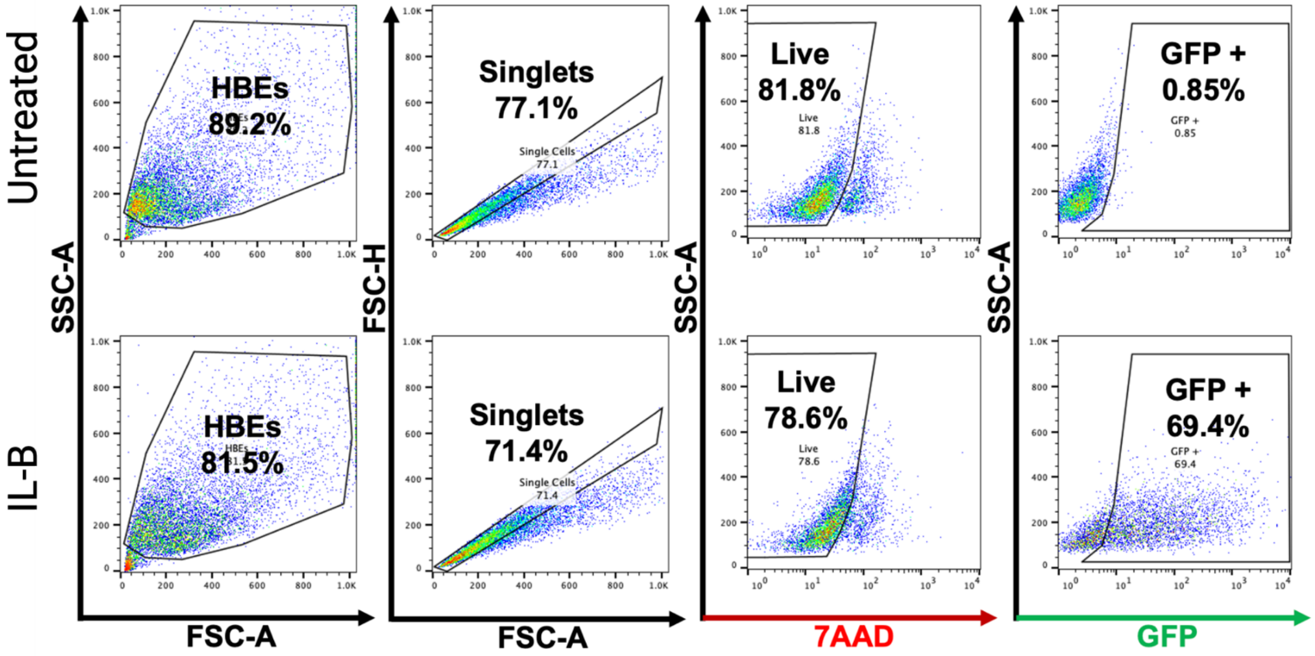


**Figure S1.** Representative flow cytometry plots depicting gating strategy for all lipid nanoparticle (LNP) screening experiments delivering mRNA encoding green fluorescent protein (GFP).

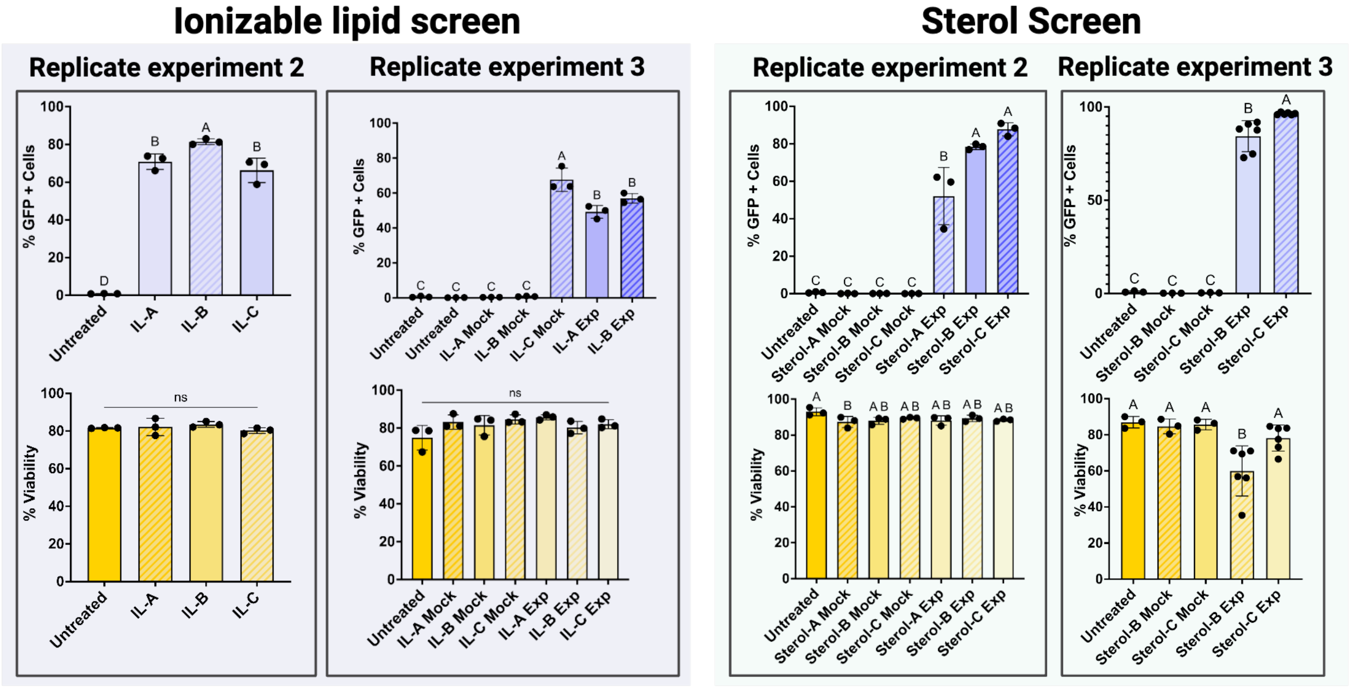


**Figure S2.** Flow cytometry data depicting (top) green fluorescent protein (GFP) expression and (bottom) cell viability in 16HBE14o- cells 24 h after treatment with lipid nanoparticles (LNPs) containing various (left) ionizable lipids and (right) sterols. Cell viability was determined with 7-AAD staining. Data represent second and third biological replicates of LNP optimization studies reported in the text (Fig. 2B, 2C). One-way analyses of variance (ANOVA) were performed, with the threshold of statistical significance set at $p < 0.05$.

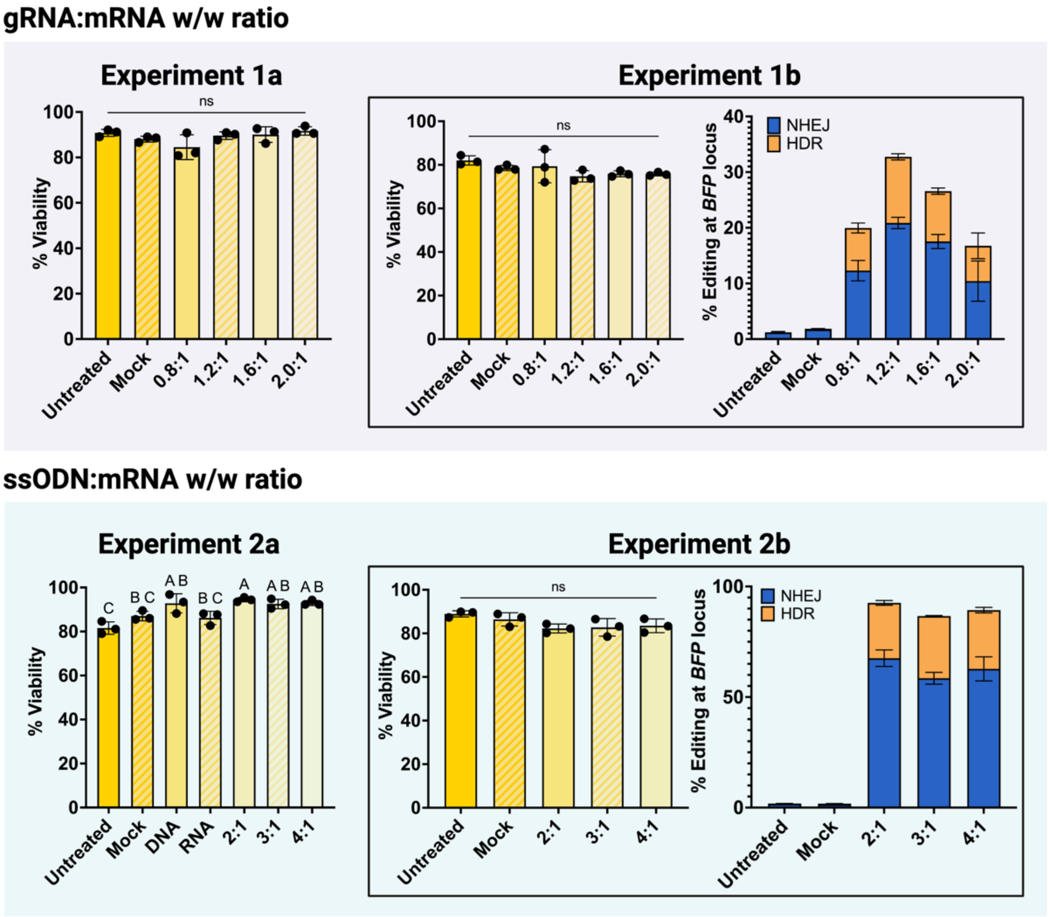


**Figure S3.** Viability and editing of BFP-16HBE14o- cells treated with LNPs loaded with Cas9 mRNA, sgRNA targeting the BFP locus, and ssODNs configured to convert BFP to GFP, at various sgRNA:mRNA w/w ratios (top) or ssODN:mRNA ratios (bottom). Cells were stained with 7-AAD viability dye and live cells were quantified *via* flow cytometry. "Experiment 1a" and "Experiment 2a" refer to the editing data reported in the text. "Experiment 1b" and "Experiment 2b" are biological replicates of the data reported in the text (Fig. 3B, 3C). One-way analyses of variance (ANOVA) were performed, with the threshold of statistical significance set at $p < 0.05$.

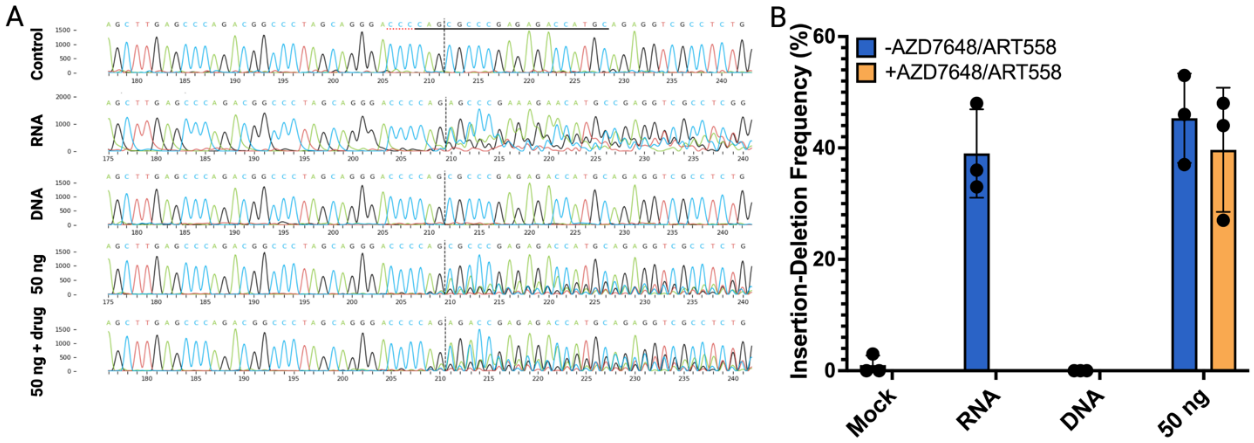


**Figure S4.** (**A**) Traces representing analysis of insertion-deletions (indels) producing allelic disruption of the 5'UTR within the endogenous cystic fibrosis transmembrane conductance regulator (*CFTR*) gene and (**B**) plots summarizing indel frequency in 16HBEgeG542x cells treated with lipid nanoparticles (LNPs). "RNA" samples were treated with LNPs loaded with Cas9 mRNA and single guide RNA (sgRNA) only, "DNA" samples were treated with LNPs loaded with a linear double-stranded DNA (ldsDNA) donor cassette encoding *CFTR,* and "50 ng" samples were treated with LNPs loaded with Cas9 mRNA, sgRNA, and ldsDNA all in one formulation at a dose of 50 ng mRNA. Samples labeled "+ drug" or "+AZD7648/ART588" were co-treated with 0.2 µM AZD-7648 and 0.5 µM ART-558. Analysis was conducted using Synthego's online Inference of CRISPR Edits (ICE) tool.

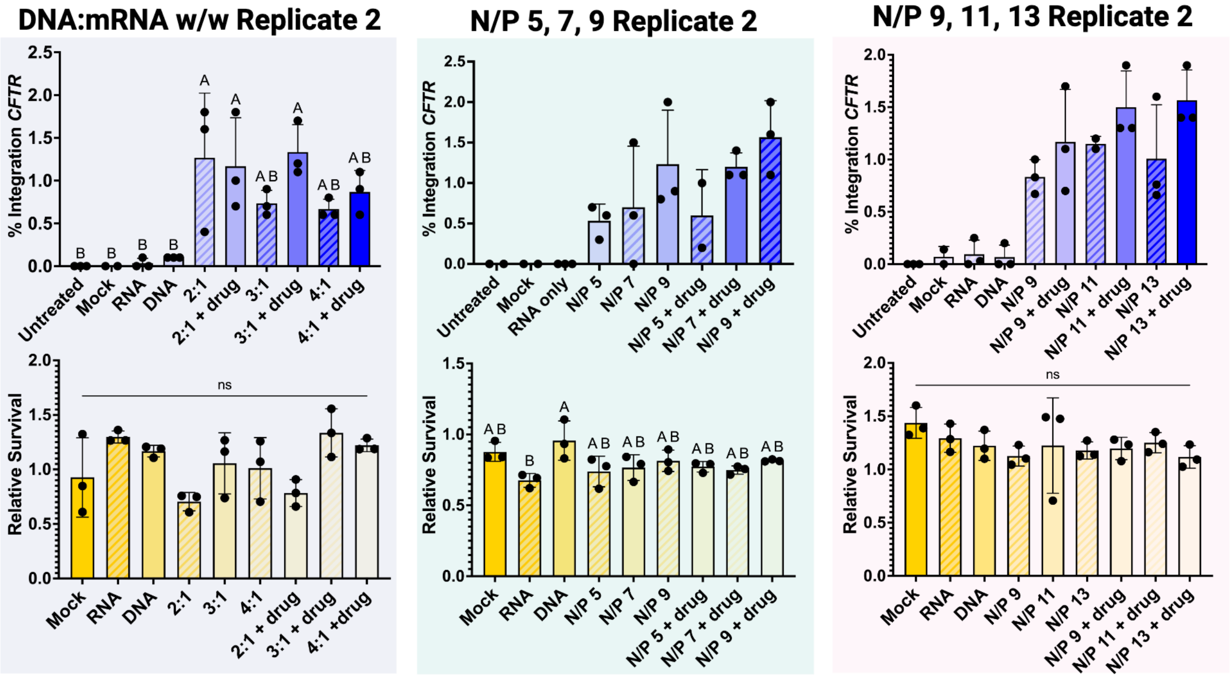


**Figure S5.** (Top) Droplet digital PCR (ddPCR) data depicting rates of integration of linear double-stranded DNA (ldsDNA) donor cassettes and (bottom) metabolic tetrazolium salt (MTS) assay data depicting 24 h cell survival in 16HBEgeG542x cells after treatment with lipid nanoparticles (LNPs) loaded with mRNA encoding Cas9, single guide RNA (sgRNA) targeting the endogenous CFTR 5'UTR, and ldsDNA encoding CFTR. Data are from biological replicates of experimental data reported in the text (Fig. 6B-G). One-way analyses of variance (ANOVA) were performed on normally distributed data, with the threshold of statistical significance set at $p < 0.05$. Letters A-B represent a compact letter display of statistics wherein differences between groups labeled with the same letter are not statistically significant.

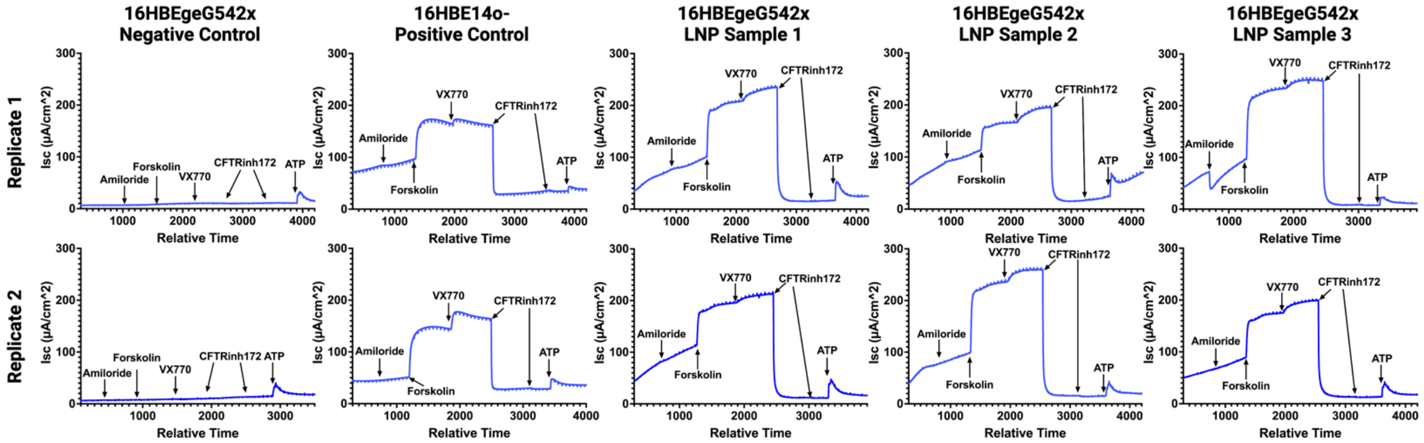


**Figure S6.** Transepithelial chloride current traces of 16HBE samples grown on SnapWell inserts and assayed in an Ussing chamber. Lipid nanoparticle (LNP) samples were edited with LNPs loaded with mRNA encoding Cas9, single guide RNA (sgRNA) targeting the endogenous *CFTR* 5'UTR, and linear double stranded DNA (ldsDNA) encoding *CFTR*. Replicates 1 & 2 represent technical replicates.

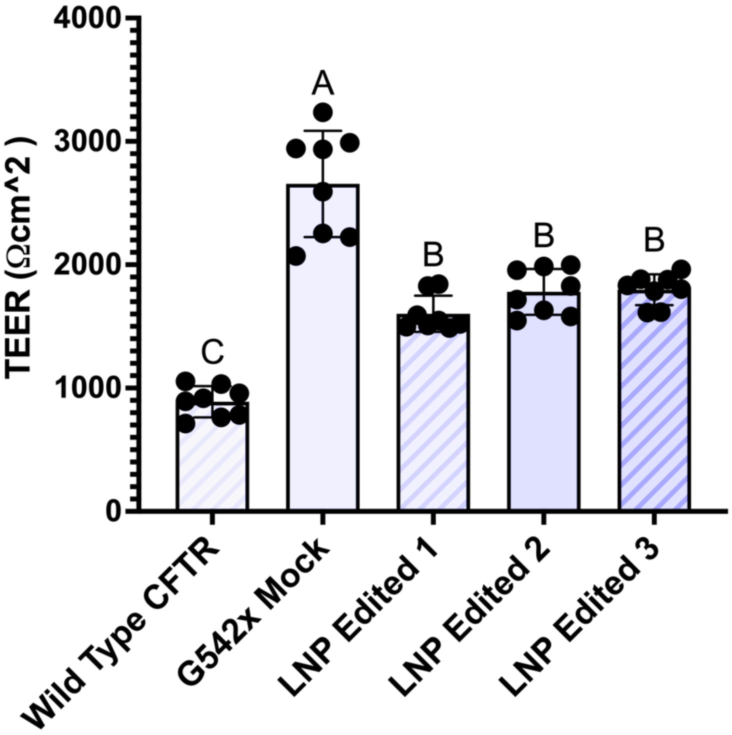


**Figure S7.** Transepithelial electrical resistance (TEER) values from transepithelial current clamp (TECC) assays of 16HBEG542x cells bulk-edited with LNPs to achieve *ca.* 3% integration of *CFTR.* 16HBE14o- cells were used as a positive control and unedited 16HBEgeG542x were used as a negative control. No significant difference was found between LNP-edited biological replicates. A one-way analysis of variance (ANOVA) was performed, with the threshold of statistical significance set at $p < 0.05$.

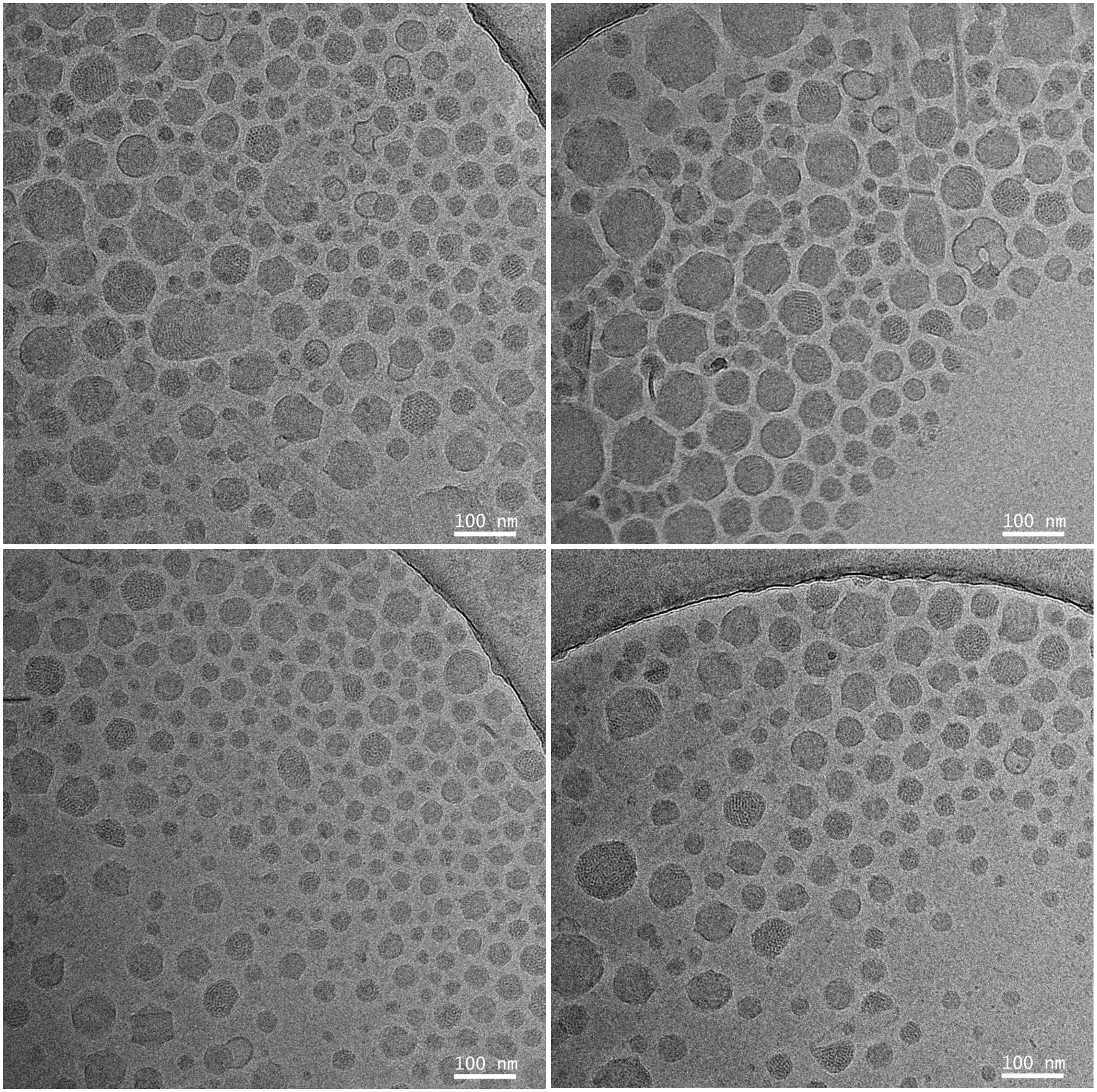


**Figure S8.** Cryogenic transmission electron microscopy (cryo-TEM) images depicting lipid nanoparticles (LNPs) loaded with mRNA transcripts encoding Cas9 and a single guide RNA (sgRNA) targeting the 5'UTR of the cystic fibrosis transmembrane conductance regulator (*CFTR*) gene.

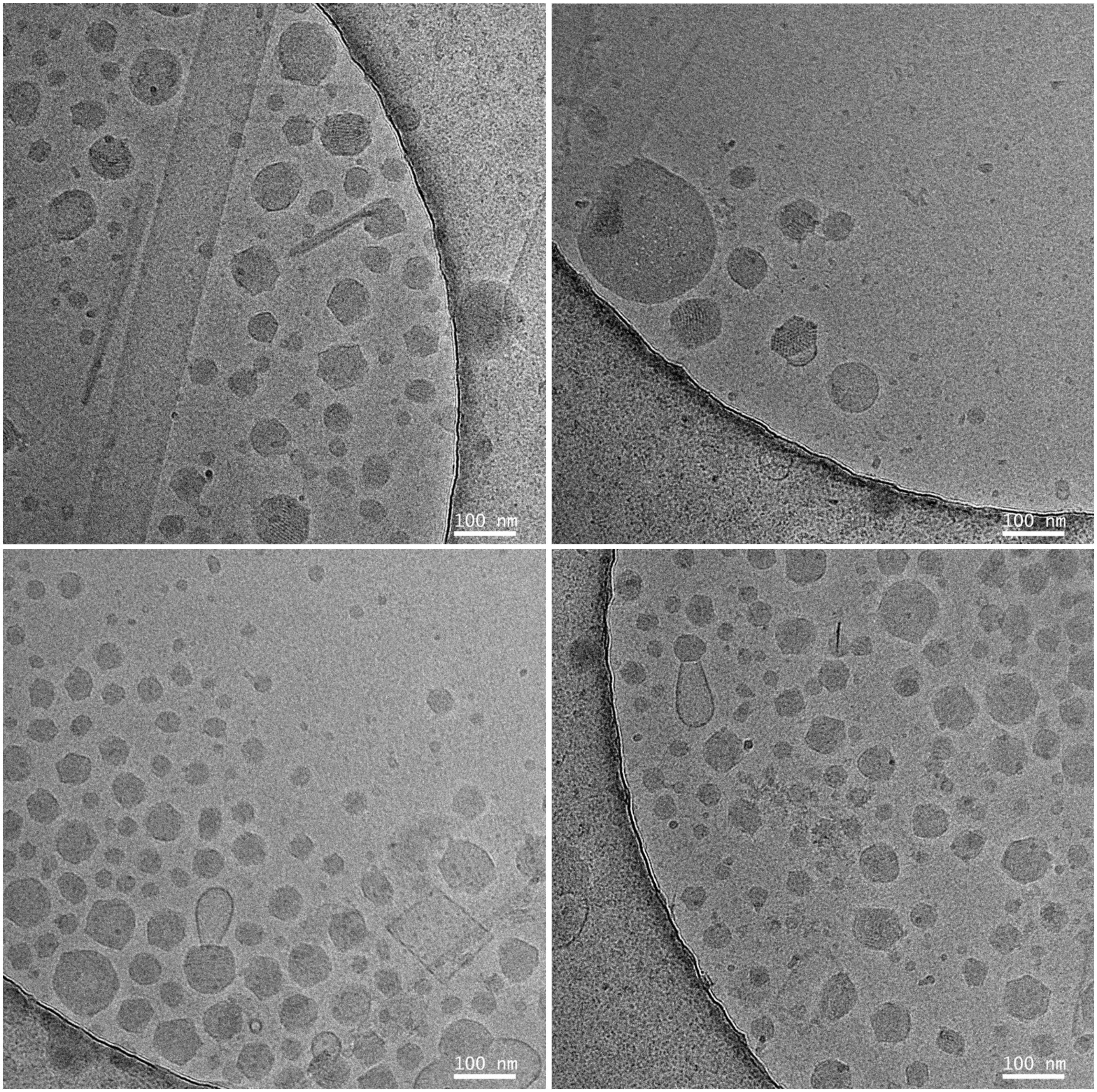


**Figure S9.** Cryogenic transmission electron microscopy (cryo-TEM) images depicting lipid nanoparticles (LNPs) loaded with linear double-stranded DNA (ldsDNA) encoding the cystic fibrosis transmembrane conductance regulator (*CFTR*) gene.

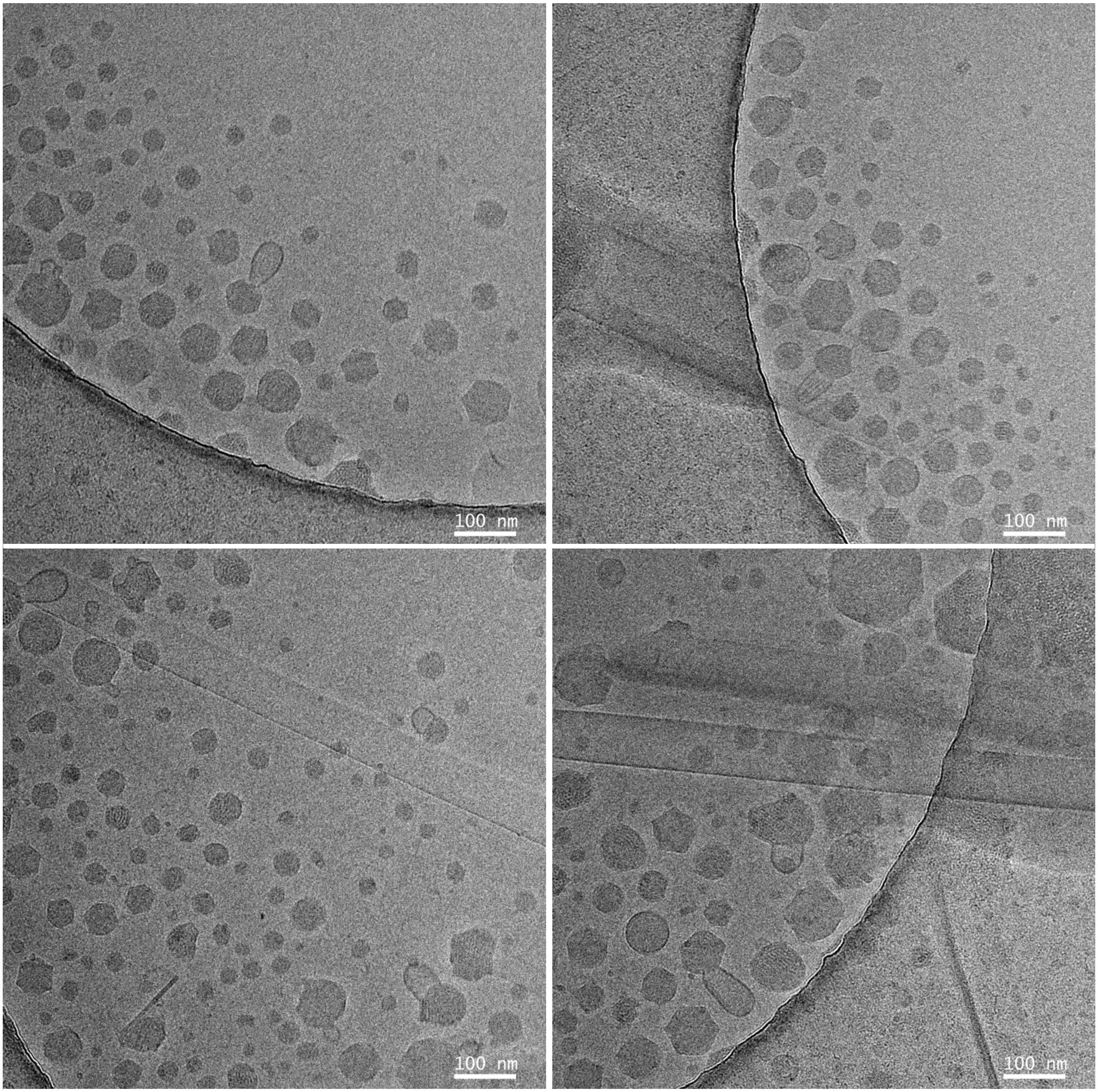


**Figure S10.** Cryogenic transmission electron microscopy (cryo-TEM) images depicting lipid nanoparticles (LNPs) loaded with mRNA transcripts encoding Cas9, a single guide RNA (sgRNA) targeting the 5'UTR of the cystic fibrosis transmembrane conductance regulator (*CFTR*) gene, and linear double-stranded DNA (ldsDNA) *CFTR* gene.

### Supplementary References

1. H. C. Valley, K. M. Bukis, A. Bell, Y. Cheng, E. Wong, N. J. Jordan, N. E. Allaire, A. Sivachenko, F. Liang, H. Bihler, P. J. Thomas, J. Mahiou, M. Mense, Isogenic cell models of cystic fibrosis-causing variants in natively expressing pulmonary epithelial cells. *J Cyst Fibros* **18**, 476-483 (2019).
